## Supplementary material for "Factors enforcing the species boundary between the human pathogens *Cryptococcus neoformans* and *Cryptococcus deneoformans*": S1 Table

**S1 Table. (A) Frequency of bald basidia produced by hybrid genetic crosses, and (B) one-way ANOVA and Tukey’s HSD post hoc statistical tests for frequencies of bald basidia.**

| **Genetic Cross** | **Mating Patch** | **Bald Basidia** | **Total Basidia** | **Frequency** |
| --- | --- | --- | --- | --- |
| H99α x JEC20**a** | 1 | 50 | 133 | 38% |
|  | 2 | 43 | 140 | 31% |
| KN99α *msh2*Δ x JEC20**a** | 1 | 59 | 165 | 36% |
|  | 2 | 50 | 157 | 32% |
| H99α x JEC20**a** *msh2*Δ*-1* | 1 | 50 | 155 | 32% |
|  | 2 | 50 | 137 | 37% |
| H99α x JEC20**a** *msh2*Δ*-2* | 1 | 38 | 141 | 27% |
|  | 2 | 43 | 133 | 32% |
| H99α x JEC20**a** *msh2*Δ*-3* | 1 | 63 | 165 | 38% |
|  | 2 | 47 | 134 | 35% |
| H99α x JEC20**a** *msh2*Δ*-4* | 1 | 48 | 152 | 32% |
|  | 2 | 42 | 151 | 28% |
| KN99α *msh2*Δ x JEC20**a** *msh2*Δ*-1* | 1 | 107 | 168 | 64% |
|  | 2 | 92 | 135 | 68% |
| KN99α *msh2*Δ x JEC20**a** *msh2*Δ*-2* | 1 | 122 | 187 | 65% |
|  | 2 | 120 | 190 | 63% |
| KN99α *msh2*Δ x JEC20**a** *msh2*Δ*-3* | 1 | 98 | 162 | 61% |
|  | 2 | 107 | 163 | 66% |
| KN99α *msh2*Δ x JEC20**a** *msh2*Δ*-4* | 1 | 99 | 177 | 56% |
|  | 2 | 112 | 190 | 59% |

**(B)**

**One-way ANOVA**

|  | **Degrees of Freedom** | **Sum of Squares** | **Mean Square** | **F value** | **p-Value** |
| --- | --- | --- | --- | --- | --- |
| Cross type | 2 | 0.4210 | 0.21052 | 141.5 | **2.53e-11***** |
| Residuals | 17 | 0.0253 | 0.00149 |  |  |

**Tukey’s HSD**

|  | **Difference** | **95% Confidence Interval** | | **p-Value** |
| --- | --- | --- | --- | --- |
|  |  | **Lower** | **Upper** |  |
| wildtype-unilateral | 0.01325109 | -0.06339573 | 0.0898979 | 0.8978853 |
| bilateral-unilateral | 0.29827815 | 0.25134175 | 0.3452145 | **0.0000000***** |
| bilateral-wildtype | 0.28502706 | 0.20679974 | 0.3632544 | **0.0000001***** |
