## Supplementary material for "Factors enforcing the species boundary between the human pathogens *Cryptococcus neoformans* and *Cryptococcus deneoformans*": S2 Table

**S2 Table. (A) Germination frequencies of progeny derived from hybrid and intra-specific crosses, and one-way ANOVA with Tukey’s HSD post hoc statistical tests for germination frequencies of (B) *C. neoformans* x *C. deneoformans* hybrid progeny, (C) progeny from *C. neoformans* x *C. neoformans* intraspecific crosses, and (D) *C. deneoformans* x *C. deneoformans* intraspecific crosses.**

**(A)**

| **Genetic Cross** | **Spores Dissected** | **Spores Germinated** | **Germination Frequency** |
| --- | --- | --- | --- |
| H99α x JEC20**a** | 100 | 6 | 6% |
|  | 71 | 0 | 0% |
|  | 83 | 2 | 2% |
|  | 60 | 9 | 15% |
| KN99α *msh2*Δ x JEC20**a** | 100 | 7 | 7% |
|  | 76 | 0 | 0% |
|  | 106 | 16 | 15% |
| H99α x JEC20**a** *msh2*Δ*-1* | 100 | 7 | 7% |
|  | 69 | 4 | 6% |
|  | 37 | 0 | 0% |
|  | 139 | 28 | 20% |
| H99α x JEC20**a** *msh2*Δ*-2* | 94 | 2 | 2% |
| H99α x JEC20**a** *msh2*Δ*-3* | 94 | 10 | 11% |
| H99α x JEC20**a** *msh2*Δ*-4* | 94 | 7 | 7% |
| KN99α *msh2*Δ x JEC20**a** *msh2*Δ*-1* | 100 | 24 | 24% |
|  | 63 | 26 | 41% |
|  | 42 | 13 | 31% |
|  | 86 | 30 | 35% |
| KN99α *msh2*Δ x JEC20**a** *msh2*Δ*-2* | 94 | 27 | 29% |
| KN99α *msh2*Δ x JEC20**a** *msh2*Δ*-3* | 94 | 24 | 26% |
| KN99α *msh2*Δ x JEC20**a** *msh2*Δ*-4* | 94 | 17 | 18% |
| H99α x KN99**a** | 75 | 62 | 83% |
|  | 76 | 69 | 91% |
|  | 77 | 57 | 74% |
| KN99α *msh2*Δ x KN99**a** | 57 | 49 | 86% |
|  | 73 | 63 | 86% |
|  | 80 | 68 | 85% |
| KN99α *msh2*Δ x KN99**a** *msh2*Δ*-1* | 73 | 47 | 64% |
|  | 76 | 56 | 74% |
|  | 74 | 49 | 66% |
| KN99α *msh2*Δ x KN99**a** *msh2*Δ*-2* | 74 | 59 | 80% |
|  | 76 | 53 | 70% |
|  | 75 | 54 | 72% |
| JEC21α x JEC20**a** | 78 | 65 | 83% |
|  | 78 | 56 | 72% |
| JEC21α x JEC20**a** *msh2*Δ*-1* | 42 | 32 | 76% |
|  | 68 | 58 | 85% |
|  | 47 | 34 | 72% |
| JEC21α *msh2*Δ*-1* x JEC20**a** *msh2*Δ*-1* | 78 | 26 | 33% |
|  | 78 | 34 | 44% |
| JEC21α *msh2*Δ*-2* x JEC20**a** *msh2*Δ*-1* | 78 | 27 | 35% |
|  | 78 | 26 | 33% |

**(B)**

**One-way ANOVA**

|  | **Degrees of Freedom** | **Sum of Squares** | **Mean Square** | **F value** | **p-Value** |
| --- | --- | --- | --- | --- | --- |
| Cross type | 2 | 2296.8 | 1148.4 | 24.55 | **7.2e-06***** |
| Residuals | 18 | 842.1 | 46.8 |  |  |

**Tukey’s HSD**

|  | **Difference** | **95% Confidence Interval** | | **p-Value** |
| --- | --- | --- | --- | --- |
|  |  | **Lower** | **Upper** |  |
| wildtype-unilateral | -1.75000 | -12.07737 | 8.577373 | 0.9025916 |
| bilateral-unilateral | 21.64286 | 13.04023 | 30.245488 | **0.0000138***** |
| bilateral-wildtype | 23.39286 | 12.45145 | 34.334264 | **0.0000990***** |

**(C)**

**One-way ANOVA**

|  | **Degrees of Freedom** | **Sum of Squares** | **Mean Square** | **F value** | **p-Value** |
| --- | --- | --- | --- | --- | --- |
| Cross type | 2 | 536.0 | 268.00 | 8.201 | **0.00938**** |
| Residuals | 9 | 294.1 | 32.68 |  |  |

**Tukey’s HSD**

|  | **Difference** | **95% Confidence Interval** | | **p-Value** |
| --- | --- | --- | --- | --- |
|  |  | **Lower** | **Upper** |  |
| wildtype-unilateral | -3.261389 | -16.29352 | 9.770739 | 0.7701868 |
| bilateral-unilateral | -14.797001 | -26.08316 | -3.510846 | **0.0130667*** |
| bilateral-wildtype | -11.535611 | -22.82177 | -0.249457 | **0.0454110*** |

**(D)**

**One-way ANOVA**

|  | **Degrees of Freedom** | **Sum of Squares** | **Mean Square** | **F value** | **p-Value** |
| --- | --- | --- | --- | --- | --- |
| Cross type | 2 | 3841 | 1920.4 | 50.4 | **0.000177***** |
| Residuals | 6 | 229 | 38.1 |  |  |

**Tukey’s HSD**

|  | **Difference** | **95% Confidence Interval** | | **p-Value** |
| --- | --- | --- | --- | --- |
|  |  | **Lower** | **Upper** |  |
| wildtype-unilateral | -0.3775706 | -17.66720 | 16.1206 | 0.9975288 |
| bilateral-unilateral | -41.7237244 | -56.18927 | -27.25818 | **0.0002844***** |
| bilateral-wildtype | -41.3461538 | -57.74854 | -24.94377 | **0.0006005***** |
