## Supplementary material for "Factors enforcing the species boundary between the human pathogens *Cryptococcus neoformans* and *Cryptococcus deneoformans*": S3 Table

**S3 Table. (A) Self-filamentation on YPD medium in hybrid and intraspecific progeny, and (B) one-way ANOVA statistical test for self-filamentation of hybrid progeny.**

| **Genetic Cross** | **Progeny Assessed** | **Self-filamentous Progeny on YPD** | **Frequency** |
| --- | --- | --- | --- |
| H99α x JEC20**a** | 9 | 3 | 33% |
| H99α x JEC20**a** | 6 | 1 | 17% |
| KN99α *msh2*Δ x JEC20**a** | 23 | 8 | 35% |
| H99α x JEC20**a** *msh2*Δ*-1* | 37 | 3 | 8% |
| H99α x JEC20**a** *msh2*Δ*-2* | 2 | 1 | 50% |
| H99α x JEC20**a** *msh2*Δ*-3* | 10 | 0 | 0% |
| H99α x JEC20**a** *msh2*Δ*-4* | 7 | 0 | 0% |
| KN99α *msh2*Δ x JEC20**a** *msh2*Δ*-1* | 96 | 44 | 46% |
| KN99α *msh2*Δ x JEC20**a** *msh2*Δ*-2* | 27 | 15 | 56% |
| KN99α *msh2*Δ x JEC20**a** *msh2*Δ*-3* | 24 | 12 | 50% |
| KN99α *msh2*Δ x JEC20**a** *msh2*Δ*-4* | 17 | 5 | 29% |
| H99α x KN99**a** | 24 | 0 | 0% |
| KN99α *msh2*Δ x KN99**a** | 24 | 0 | 0% |
| KN99α *msh2*Δ x KN99**a** *msh2*Δ*-1* | 24 | 0 | 0% |
| KN99α *msh2*Δ x KN99**a** *msh2*Δ*-2* | 24 | 0 | 0% |
| JEC21α x JEC20**a** | 24 | 0 | 0% |
| JEC21α x JEC20**a** *msh2*Δ*-1* | 24 | 0 | 0% |
| JEC21α *msh2*Δ x JEC20**a** *msh2*Δ*-1* | 24 | 0 | 0% |
| JEC21α *msh2*Δ x JEC20**a** *msh2*Δ*-2* | 24 | 0 | 0% |

**(B)**

**One-way ANOVA**

|  | **Degrees of Freedom** | **Sum of Squares** | **Mean Square** | **F value** | **p-Value** |
| --- | --- | --- | --- | --- | --- |
| Cross type | 2 | 1623 | 811.4 | 2.527 | 0.141 |
| Residuals | 8 | 2569 | 321.1 |  |  |
