## Supplementary material for "Factors enforcing the species boundary between the human pathogens *Cryptococcus neoformans* and *Cryptococcus deneoformans*": S5 Table

**S5 Table.** **(A) Heterozygosity level of the hybrid progeny, and (B) Kruskal-Wallis and Dunn statistical tests.**

**(A)**

| **Cross** | **Progeny** | **HET sites** | **HOM-VAR sites** | **Total Sites** | **HET %** |
| --- | --- | --- | --- | --- | --- |
| H99α X JEC20**a** | YX1 | 76,512 | 434,032 | 510,544 | 14.99 |
|  | YX3 | 238,400 | 172,080 | 410,480 | 58.08 |
|  | YX4 | 710,400 | 94,481 | 804,881 | 88.26 |
|  | YX5 | 689,492 | 94,899 | 784,391 | 87.9 |
|  | YX6 | 282,507 | 523,391 | 805,898 | 35.05 |
|  | YX7 | 715,778 | 95,073 | 810,851 | 88.27 |
|  | YX8 | 957,100 | 40,338 | 997,438 | 95.96 |
| H99α x JEC20**a** *msh2*Δ*-1* | YX10 | 834,476 | 183,602 | 1,018,078 | 81.97 |
|  | YX12 | 553,150 | 192,686 | 745,836 | 74.17 |
|  | YX14 | 885,929 | 89,431 | 975,360 | 90.83 |
|  | YX16 | 523,113 | 209,443 | 732,556 | 71.41 |
|  | YX17 | 831,142 | 9,071 | 840,213 | 98.92 |
|  | YX19 | 798,948 | 65,962 | 864,910 | 92.37 |
| KN99α *msh2*Δ*-1* x JEC20**a** | YX43 | 830,792 | 108,761 | 939,553 | 88.42 |
|  | YX44 | 1,000,885 | 3,211 | 1,004,096 | 99.68 |
|  | YX45 | 913,322 | 58,386 | 971,708 | 93.99 |
|  | YX47 | 996,266 | 3,188 | 999,454 | 99.68 |
|  | YX48 | 904,976 | 59,593 | 964,569 | 93.82 |
|  | YX49 | 871,725 | 135,549 | 1,007,274 | 86.54 |
| KN99α *msh2*Δ*-1* x JEC20**a** *msh2*Δ*-1* | YX56 | 853,776 | 40,830 | 894,606 | 95.44 |
|  | YX58 | 949,380 | 2,756 | 952,136 | 99.71 |
|  | YX59 | 994,854 | 2,791 | 997,645 | 99.72 |
|  | YX60 | 920,161 | 2,829 | 922,990 | 99.69 |
|  | YX61 | 778,101 | 184,215 | 962,316 | 80.86 |
|  | YX64 | 839,153 | 41,706 | 880,859 | 95.27 |
|  | YX65 | 995,355 | 2,790 | 998,145 | 99.72 |
|  | YX70 | 852,332 | 41,396 | 893,728 | 95.37 |

**(B)**

**Wilcoxon / Kruskal-Wallis Test**

| **Level** | **Count** | **Score Sum** | **Exp. Score** | **Score Mean** | **(Mean-Mean0)/Std0** |
| --- | --- | --- | --- | --- | --- |
| H99α X JEC20**a** | 7 | 56.000 | 98.000 | 8.0000 | -2.297 |
| H99α x JEC20**a** *msh2*Δ-1 | 6 | 64.000 | 84.000 | 10.6667 | -1.138 |
| KN99α *msh2*Δ*-1* x JEC20**a** | 6 | 96.000 | 84.000 | 16.0000 | 0.671 |
| KN99α *msh2*Δ*-1* x JEC20**a** *msh2*Δ*-1* | 8 | 162.000 | 112.000 | 20.2500 | 2.629 |

**1-Way Test, Chi-square Approximation**

| ChiSquare | DF | Prob>ChiSq |
| --- | --- | --- |
| 10.4058 | 3 | **0.0154*** |

**Nonparametric comparisons for all pairs using Dunn method**

| **Level** | **Level** | **Score Mean Dif** | **Std Err Dif** | **Z** | **p-Value** |
| --- | --- | --- | --- | --- | --- |
| KN99α *msh2*Δ*-1* x JEC20**a** *msh2*Δ*-1* | H99α x JEC20**a** | 12.11607 | 4.106665 | 2.950343 | **0.0190*** |
| KN99α *msh2*Δ*-1* x JEC20**a** *msh2*Δ*-1* | H99α x JEC20**a** *msh2*Δ-1 | 9.43750 | 4.285298 | 2.202297 | 0.1659 |
| KN99α *msh2*Δ*-1* x JEC20**a** | H99α x JEC20**a** | 7.84524 | 4.414532 | 1.777139 | 0.4533 |
| KN99α *msh2*Δ*-1* x JEC20**a** | H99α x JEC20**a** *msh2*Δ-1 | 5.16667 | 4.581177 | 1.127803 | 1.0000 |
| KN99α *msh2*Δ*-1* x JEC20**a** *msh2*Δ*-1* | KN99α *msh2*Δ*-1* x JEC20**a** | 4.10417 | 4.285298 | 0.957732 | 1.0000 |
| H99α x JEC20**a** *msh2*Δ*-1* | H99α x JEC20**a** | 2.51190 | 4.414532 | 0.569008 | 1.0000 |
