## Supplementary material for "Factors enforcing the species boundary between the human pathogens *Cryptococcus neoformans* and *Cryptococcus deneoformans*": S7 Table

**S7 Table. Genomic locations of telomeric repeats added *de novo* at chromosome breaks in progeny derived from unilateral and bilateral hybrid crosses of *C. neoformans* × *C. deneoformans* *msh2* mutants, and in a single progeny derived from a bilateral intraspecific cross of *C. neoformans* *msh2* mutants.**

| **Progeny** | **Genetic cross** | **Ref genome,**  **Chr. no.** | **Chr. location**  (coordinates in bp) | **Genomic features** | **Figure** |
| --- | --- | --- | --- | --- | --- |
| YX56 | KN99α *msh2*Δ × JEC20**a** *msh2*Δ*-1* | H99, Chr3 | 5'end (16,484) | gene (CNAG_06983) | S11D Fig. |
| YX64 | KN99α *msh2*Δ × JEC20**a** *msh2*Δ*-1* | H99, Chr3 | 5'end (16,484) | gene (CNAG_06983) | S11D Fig. |
| YX70* | KN99α *msh2*Δ × JEC20**a** *msh2*Δ*-1* | H99, Chr3 | 5'end (16,484) | gene (CNAG_06983) | S11D Fig. |
| YX43 | KN99α *msh2*Δ × JEC20**a** | H99, Chr10 | 5’end (6,479) | 3'UTR (CNAG_07836) | S11E Fig. |
| YX44 | KN99α *msh2* Δ × JEC20**a** | H99, Chr10 | 5’end (6,479) | 3’UTR (CNAG_07836) | S11E Fig. |
| YX45* | KN99α *msh2* Δ × JEC20**a** | H99, Chr10 | 5’end (6,479) | 3’UTR (CNAG_07836) | S11E Fig. |
| YX46 | KN99α *msh2*Δ × JEC20**a** | H99, Chr10 | 5’end (6,479) | 3’UTR (CNAG_07836) | S11E Fig. |
| YX47 | KN99α *msh2*Δ × JEC20**a** | H99, Chr10 | 5’end (6,479) | 3’UTR (CNAG_07836) | S11E Fig. |
| YX48* | KN99α *msh2*Δ × JEC20**a** | H99, Chr10 | 5’end (6,479) | 3’UTR (CNAG_07836) | S11E Fig. |
| YX49 | KN99α *msh2*Δ × JEC20**a** | H99, Chr10 | 5’end (6,479) | 3’UTR (CNAG_07836) | S11E Fig. |
| YX61 | KN99α *msh2*Δ × JEC20**a** *msh2*Δ*-1* | H99, Chr13 | q-arm (184,59) | intergenic | S11F Fig. |
| YX58* | KN99α *msh2*Δ × JEC20**a** *msh2*Δ*-1* | JEC21, Chr6 | 3’end (1,405,933) | T1 DNA transposon | S11C Fig. |
| YX70* | KN99α *msh2*Δ × JEC20**a** *msh2*Δ*-1* | JEC21, Chr6 | 3’end (1,405,933) | T1 DNA transposon | S11C Fig. |
| YX45* | KN99α *msh2*Δ × JEC20**a** | JEC21, Chr14 | centromere 14 (630,636) | TCN2 LTR | S11A Fig. |
| YX48* | KN99α *msh2*Δ × JEC20**a** | JEC21, Chr14 | centromere 14 (630,636) | TCN2 LTR | S11A Fig. |
| YX58* | KN99α *msh2*Δ × JEC20**a** *msh2*Δ*-1* | JEC21, Chr9 | centromere 9 (791,053) | TCN1 LTR | S11B Fig. |
| # 3 | KN99α *msh2*Δ × KN99**a** *msh2*Δ*-2* | H99, Chr13 | 5’end 9 (17,369) | LTR11 | S12 Fig. |

*progeny in which more than one chromosome underwent breakage and healing via *de novo* telomere addition.
