## Supplementary material for "Factors enforcing the species boundary between the human pathogens *Cryptococcus neoformans* and *Cryptococcus deneoformans*": S8 Table

**S8 Table. Strains used in this study.**

| **Strain** | **Genotype** | **Comment** | **Source** |
| --- | --- | --- | --- |
| H99α | *MAT*α | H99F isolate | [34] |
| JEC20**a** | *MAT***a** | Also known as B-4476 | [70] |
| JEC21α | *MAT*α | Also known as B-4500 | [70] |
| *msh2*Δ | *msh2*Δ::*NAT, MAT*α | From 2015 Madhani deletion collection; KN99α genetic background | [43] |
| *fur1*Δ | *fur1*Δ::*NAT, MAT*α | From 2015 Madhani deletion collection; KN99α genetic background | [43] |
| YX2 | *msh2*Δ*-1*::*NAT, MAT***a** | Independent mutant *msh2*Δ*-1*; JEC20 genetic background | This study |
| SJP537 | *msh2*Δ*-2*::*NAT, MAT***a** | Independent mutant *msh2*Δ*-2*; JEC20 genetic background | This study |
| SJP538 | *msh2*Δ*-3*::*NAT, MAT***a** | Independent mutant *msh2*Δ*-3*; JEC20 genetic background | This study |
| SJP540 | *msh2*Δ*-4*::*NAT, MAT***a** | Independent mutant *msh2*Δ*-4*; JEC20 genetic background | This study |
| YX1 | *C. neoformans MAT*α,  *C. deneoformans* *MAT***a** | Hybrid progeny from wildtype H99α x JEC20**a** cross | This study |
| YX3 | *C. neoformans MAT*α, *C. deneoformans* *MAT***a** | Hybrid progeny from wildtype H99α x JEC20**a** cross | This study |
| YX4 | *C. neoformans MAT*α, *C. deneoformans* *MAT***a** | Hybrid progeny from wildtype H99α x JEC20**a** cross | This study |
| YX5 | *C. neoformans MAT*α, *C. deneoformans* *MAT***a** | Hybrid progeny from wildtype H99α x JEC20**a** cross | This study |
| YX6 | *C. neoformans MAT*α | Hybrid progeny from wildtype H99α x JEC20**a** cross | This study |
| YX7 | *C. neoformans MAT*α, *C. deneoformans* *MAT***a** | Hybrid progeny from wildtype H99α x JEC20**a** cross | This study |
| YX8 | *C. neoformans MAT*α, *C. deneoformans* *MAT***a** | Hybrid progeny from wildtype H99α x JEC20**a** cross | This study |
| YX10 | *C. neoformans MAT*α, *C. deneoformans* *MAT***a** | Hybrid progeny from wildtype H99α x JEC20**a** cross | This study |
| YX12 | *C. neoformans MAT*α, *C. deneoformans* *MAT***a** | Hybrid progeny from KN99α *msh2*Δ x JEC20**a** cross | This study |
| YX14 | *C. neoformans MAT*α, *C. deneoformans* *MAT***a** | Hybrid progeny from KN99α *msh2*Δ x JEC20**a** cross | This study |
| YX16 | *C. neoformans MAT*α, *C. deneoformans* *MAT***a** | Hybrid progeny from KN99α *msh2*Δ x JEC20**a** cross | This study |
| YX17 | *C. neoformans MAT*α, *C. deneoformans* *MAT***a** | Hybrid progeny from KN99α *msh2*Δ x JEC20**a** cross | This study |
| YX19 | *C. neoformans MAT*α, *C. deneoformans* *MAT***a** | Hybrid progeny from KN99α *msh2*Δ x JEC20**a** cross | This study |
| YX43 | *C. neoformans MAT*α, *C. deneoformans* *MAT***a** | Hybrid progeny from H99α x JEC20**a** *msh2*Δ*-1* cross | This study |
| YX44 | *C. neoformans MAT*α, *C. deneoformans* *MAT***a** | Hybrid progeny from H99α x JEC20**a** *msh2*Δ*-1* cross | This study |
| YX45 | *C. neoformans MAT*α, *C. deneoformans* *MAT***a** | Hybrid progeny from H99α x JEC20**a** *msh2*Δ*-1* cross | This study |
| YX47 | *C. neoformans MAT*α, *C. deneoformans* *MAT***a** | Hybrid progeny from H99α x JEC20**a** *msh2*Δ*-1* cross | This study |
| YX48 | *C. neoformans MAT*α, *C. deneoformans* *MAT***a** | Hybrid progeny from H99α x JEC20**a** *msh2*Δ*-1* cross | This study |
| YX49 | *C. neoformans MAT*α, *C. deneoformans* *MAT***a** | Hybrid progeny from H99α x JEC20**a** *msh2*Δ*-1* cross | This study |
| YX56 | *C. neoformans MAT*α, *C. deneoformans* *MAT***a** | Hybrid progeny from KN99α *msh2*Δ x JEC20**a** *msh2*Δ*-1* cross | This study |
| YX58 | *C. neoformans MAT*α, *C. deneoformans* *MAT***a** | Hybrid progeny from KN99α *msh2*Δ x JEC20**a** *msh2*Δ*-1* cross | This study |
| YX59 | *C. neoformans MAT*α, *C. deneoformans* *MAT***a** | Hybrid progeny from KN99α *msh2*Δ x JEC20**a** *msh2*Δ*-1* cross | This study |
| YX60 | *C. neoformans MAT*α, *C. deneoformans* *MAT***a** | Hybrid progeny from KN99α *msh2*Δ x JEC20**a** *msh2*Δ*-1* cross | This study |
| YX61 | *C. neoformans MAT*α, *C. deneoformans* *MAT***a** | Hybrid progeny from KN99α *msh2*Δ x JEC20**a** *msh2*Δ*-1* cross | This study |
| YX64 | *C. neoformans MAT*α, *C. deneoformans* *MAT***a** | Hybrid progeny from KN99α *msh2*Δ x JEC20**a** *msh2*Δ*-1* cross | This study |
| YX65 | *C. neoformans MAT*α, *C. deneoformans* *MAT***a** | Hybrid progeny from KN99α *msh2*Δ x JEC20**a** *msh2*Δ*-1* cross | This study |
| YX70 | *C. neoformans MAT*α, *C. deneoformans* *MAT***a** | Hybrid progeny from KN99α *msh2*Δ x JEC20**a** *msh2*Δ*-1* cross | This study |
| AI187 | *C. neoformans ADE2*/*ade2* *URA5*/*ura5* *MAT***a**/*MAT*α | Self-filamentous diploid | [92] |
| XL143 | *C. deneoformans* *MAT*α/α diploid | Derived from JEC21 blastospore dissection | [57] |
| KN99**a** | *MAT***a** |  | [72] |
| SJP592 | *msh2*Δ*-1*::*NAT*, *MAT***a** | KN99**a** independent mutant *msh2*Δ*-1*; derived from a KN99α *msh2*Δ x KN99**a** genetic cross | This study |
| SJP593 | *msh2*Δ*-2*::*NAT*, *MAT***a** | KN99**a** independent mutant *msh2*Δ*-2*; derived from a KN99α *msh2*Δ x KN99**a** genetic cross | This study |
| SJP594 | *msh2*Δ*-1*::*NAT*, *MAT*α | JEC21α independent mutant *msh2*Δ*-1*; derived from a JEC20**a** *msh2*Δ*-1* x JEC21α genetic cross | This study |
| SJP595 | *msh2*Δ*-2*::*NAT*, *MAT*α | JEC21α independent mutant *msh2*Δ*-2*; derived from a JEC20**a** *msh2*Δ*-1* x JEC21α genetic cross | This study |
| SJP596 | *C. neoformans MAT***a** | Intra-specific progeny #1 from H99α x KN99**a** | This study |
| SJP597 | *C. neoformans MAT***a** | Intra-specific progeny #2 from H99α x KN99**a** | This study |
| SJP599 | *C. neoformans MAT***a** | Intra-specific progeny #4 from H99α x KN99**a** | This study |
| SJP600 | *C. neoformans MAT*α | Intra-specific progeny #5 from H99α x KN99**a** | This study |
| SJP601 | *C. neoformans MAT*α | Intra-specific progeny #1 from KN99α *msh2*Δ x KN99**a** | This study |
| SJP602 | *C. neoformans MAT***a** | Intra-specific progeny #2 from KN99α *msh2*Δ x KN99**a** | This study |
| SJP603 | *C. neoformans MAT***a** | Intra-specific progeny #3 from KN99α *msh2*Δ x KN99**a** | This study |
| SJP604 | *C. neoformans MAT***a** | Intra-specific progeny #4 from KN99α *msh2*Δ x KN99**a** | This study |
| SJP605 | *C. neoformans MAT*α and *MAT***a** | Intra-specific progeny #5 from KN99α *msh2*Δ x KN99**a** | This study |
| SJP606 | *C. neoformans MAT*α | Intra-specific progeny #1 from KN99α *msh2*Δ x KN99**a** *msh2*Δ*-1* | This study |
| SJP607 | *C. neoformans MAT***a** | Intra-specific progeny #2 from KN99α *msh2*Δ x KN99**a** *msh2*Δ*-1* | This study |
| SJP608 | *C. neoformans MAT***a** | Intra-specific progeny #3 from KN99α *msh2*Δ x KN99**a** *msh2*Δ*-1* | This study |
| SJP609 | *C. neoformans MAT*α | Intra-specific progeny #4 from KN99α *msh2*Δ x KN99**a** *msh2*Δ*-1* | This study |
| SJP610 | *C. neoformans MAT*α | Intra-specific progeny #5 from KN99α *msh2*Δ x KN99**a** *msh2*Δ*-1* | This study |
| SJP611 | *C. neoformans MAT*α | Intra-specific progeny #1 from KN99α *msh2*Δ x KN99**a** *msh2*Δ*-2* | This study |
| SJP612 | *C. neoformans MAT***a** | Intra-specific progeny #2 from KN99α *msh2*Δ x KN99**a** *msh2*Δ*-2* | This study |
| SJP613 | *C. neoformans MAT***a** | Intra-specific progeny #3 from KN99α *msh2*Δ x KN99**a** *msh2*Δ*-2* | This study |
| SJP614 | *C. neoformans MAT*α | Intra-specific progeny #4 from KN99α *msh2*Δ x KN99**a** *msh2*Δ*-2* | This study |
| SJP615 | *C. neoformans MAT***a** | Intra-specific progeny #5 from KN99α *msh2*Δ x KN99**a** *msh2*Δ*-2* | This study |
| SJP616 | *C. deneoformans MAT***a** | Intra-specific progeny #1 from JEC21α x JEC20**a** | This study |
| SJP617 | *C. deneoformans MAT***a** | Intra-specific progeny #2 from JEC21α x JEC20**a** | This study |
| SJP618 | *C. deneoformans MAT*α | Intra-specific progeny #3 from JEC21α x JEC20**a** | This study |
| SJP619 | *C. deneoformans MAT*α | Intra-specific progeny #4 from JEC21α x JEC20**a** | This study |
| SJP621 | *C. deneoformans MAT***a** | Intra-specific progeny #1 from JEC21α x JEC20**a** *msh2*Δ*-1* | This study |
| SJP622 | *C. deneoformans MAT*α | Intra-specific progeny #2 from JEC21α x JEC20**a** *msh2*Δ*-1* | This study |
| SJP623 | *C. deneoformans MAT*α | Intra-specific progeny #3 from JEC21α x JEC20**a** *msh2*Δ*-1* | This study |
| SJP624 | *C. deneoformans MAT*α | Intra-specific progeny #4 from JEC21α x JEC20**a** *msh2*Δ*-1* | This study |
| SJP625 | *C. deneoformans MAT*α | Intra-specific progeny #5 from JEC21α x JEC20**a** *msh2*Δ*-1* | This study |
| SJP626 | *C. deneoformans MAT***a** | Intra-specific progeny #1 from JEC21α *msh2*Δ*-1* x JEC20**a** *msh2*Δ*-1* | This study |
| SJP627 | *C. deneoformans MAT***a** | Intra-specific progeny #2 from JEC21α *msh2*Δ*-1* x JEC20**a** *msh2*Δ*-1* | This study |
| SJP628 | *C. deneoformans MAT***a** | Intra-specific progeny #3 from JEC21α *msh2*Δ*-1* x JEC20**a** *msh2*Δ*-1* | This study |
| SJP629 | *C. deneoformans MAT***a** | Intra-specific progeny #4 from JEC21α *msh2*Δ*-1* x JEC20**a** *msh2*Δ*-1* | This study |
| SJP630 | *C. deneoformans MAT*α | Intra-specific progeny #5 from JEC21α *msh2*Δ*-1* x JEC20**a** *msh2*Δ*-1* | This study |
| SJP631 | *C. deneoformans MAT***a** | Intra-specific progeny #1 from JEC21α *msh2*Δ*-2* x JEC20**a** *msh2*Δ*-1* | This study |
| SJP632 | *C. deneoformans MAT***a** | Intra-specific progeny #2 from JEC21α *msh2*Δ*-2* x JEC20**a** *msh2*Δ*-1* | This study |
| SJP633 | *C. deneoformans MAT*α | Intra-specific progeny #3 from JEC21α *msh2*Δ*-2* x JEC20**a** *msh2*Δ*-1* | This study |
| SJP634 | *C. deneoformans MAT***a** | Intra-specific progeny #4 from JEC21α *msh2*Δ*-2* x JEC20**a** *msh2*Δ*-1* | This study |
| SJP635 | *C. deneoformans MAT***a** | Intra-specific progeny #5 from JEC21α *msh2*Δ*-2* x JEC20**a** *msh2*Δ*-1* | This study |
| XL280α | *C. deneoformans MAT*α |  | [48] |
