## Supplementary material for "Factors enforcing the species boundary between the human pathogens *Cryptococcus neoformans* and *Cryptococcus deneoformans*": S9 Table

| **Primer** | **Sequence (5’ to 3’)** | **Comment** |
| --- | --- | --- |
| JOHE45551 | TGACCGGAATCTGTCTGAGG | JEC20 *MSH2* deletion, 1061 bp upstream of start, F |
| JOHE45552 | CTGGCCGTCGTTTTATTGCTTTCGCCGGGATAC | JEC20 *MSH2* deletion, 96 bp downstream of start, R |
| JOHE45553 | GTATCCCGGCGAAAGCAATAAAACGACGGCCAG | JEC20 *MSH2* deletion, Amplifies NAT marker, F |
| JOHE45554 | CTGGAAGAAAAGAGCCTGCACAGGAAACAGCTATGAC | JEC20 *MSH2* deletion, Amplifies NAT marker, R |
| JOHE45555 | GTCATAGCTGTTTCCTGTGCAGGCTCTTTTCTTCCAG | JEC20 *MSH2* deletion, 40 bp upstream of stop, F |
| JOHE45556 | TCAATCCCTTTTCCACGAAC | JEC20 *MSH2* deletion, 1040 bp downstream of stop, R |
| JOHE45559 | GATCGGGGATGCATGAGTG | JEC20 *MSH2* deletion confirmation, 1133 bp upstream of start, F |
| JOHE45560 | GCCCCTCTGGAACCATAATC | JEC20 *MSH2* deletion confirmation, 1137 bp downstream of stop, R |
| JOHE45776 | AACGACAGACCGTTCGAAGA | JEC21 *FRR1*, F |
| JOHE45777 | GTCTGCTCGCGCTTCTGTAT | JEC21 *FRR1*, R |
| JOHE45822 | CAGGTCGACATACCGCCTTA | JEC21 *MSH2* internal, F |
| JOHE45823 | ATCCTCAAGAGCTGGGTGGT | JEC21 *MSH2* internal, R |
| JOHE45868 | GGCTACGAAACAGCATCAGC | JEC21 *FRR1* internal, F |
| JOHE45869 | GCGATCTTTGCCATGTGAAT | JEC21 *FRR1* internal, R |
| JOHE51411 | CCTCAAGAGCTGGGTGGTAAC | H99 *MSH2* internal, F |
| JOHE51412 | GCCCCTAGTTCTGGAGTGTTG | H99 *MSH2* internal, R |
| JOHE51413 | ATGTATAACGGGCGATTACGC | H99 *MSH2* upstream F |
| JOHE51414 | AGATATCGGCCATCACCTCTG | H99 *MSH2* upstream R |
| JOHE39201 | CTAACTCTACTACACCTCACGGCA | *C. neoformans STE20 MAT***a** F |
| JOHE39202 | CGCACTGCAAAATAGATAAGTCTG | *C. neoformans STE20 MAT***a** R |
| JOHE39203 | GGCTGCAATCACAGCACCTTAC | *C. neoformans STE20 MAT*α F |
| JOHE39204 | CTTCATGACATCACTCCCCTAT | *C. neoformans STE20 MAT*α R |
| JOHE39205 | CACATCTCAGATGCCATTTTACCA | *C. deneoformans STE20 MAT***a** F |
| JOHE39206 | AGCTCTAAGTCATATGGGTTATAT | *C. deneoformans STE20 MAT***a** R |
| JOHE39207 | CTTAATTCACAGCACCAGCCTA | *C. deneoformans STE20 MAT*α F |
| JOHE39208 | GGTCATCACAGTCAGTCACCAC | *C. deneoformans STE20 MAT*α R |

**S9 Table. Oligonucleotides used in this study.**
