## Supplementary material for "Factors enforcing the species boundary between the human pathogens *Cryptococcus neoformans* and *Cryptococcus deneoformans*": S10 Table

**S10 Table. NCBI data submissions related to this study.**

| Sr. no. | Strain Name | Genetic Cross | Accession Number |
| --- | --- | --- | --- |
| 1. | JEC20**a** | N.A. | SAMN14647801 |
| 2. | JEC20**a** *msh2*Δ*-1* | N.A. | SAMN14647802 |
| 3. | YX1 | H99α x JEC20**a** | SAMN14647803 |
| 4. | YX3 | H99α x JEC20**a** | SAMN14647804 |
| 5. | YX4 | H99α x JEC20**a** | SAMN14647805 |
| 6. | YX5 | H99α x JEC20**a** | SAMN14647806 |
| 7. | YX6 | H99α x JEC20**a** | SAMN14647807 |
| 8. | YX7 | H99α x JEC20**a** | SAMN14647808 |
| 9. | YX8 | H99α x JEC20**a** | SAMN14647809 |
| 10 | YX10 | H99α x JEC20**a** *msh2*Δ*-1* | SAMN14647810 |
| 11. | YX12 | H99α x JEC20**a** *msh2*Δ*-1* | SAMN14647811 |
| 12. | YX14 | H99α x JEC20**a** *msh2*Δ*-1* | SAMN14647812 |
| 13. | YX16 | H99α x JEC20**a** *msh2*Δ*-1* | SAMN14647813 |
| 14. | YX17 | H99α x JEC20**a** *msh2*Δ*-1* | SAMN14647814 |
| 15. | YX19 | H99α x JEC20**a** *msh2*Δ*-1* | SAMN14647815 |
| 16. | YX43 | KN99α *msh2*Δ x JEC20**a** | SAMN14647816 |
| 17. | YX44 | KN99α *msh2*Δ x JEC20**a** | SAMN14647817 |
| 18. | YX45 | KN99α *msh2*Δ x JEC20**a** | SAMN14647818 |
| 19. | YX47 | KN99α *msh2*Δ x JEC20**a** | SAMN14647819 |
| 20. | YX48 | KN99α *msh2*Δ x JEC20**a** | SAMN14647820 |
| 21. | YX49 | KN99α *msh2*Δ x JEC20**a** | SAMN14647821 |
| 22. | YX56 | KN99α *msh2*Δ x JEC20**a** *msh2*Δ*-1* | SAMN14647822 |
| 23. | YX58 | KN99α *msh2*Δ x JEC20**a** *msh2*Δ*-1* | SAMN14647823 |
| 24. | YX59 | KN99α *msh2*Δ x JEC20**a** *msh2*Δ*-1* | SAMN14647824 |
| 25. | YX60 | KN99α *msh2*Δ x JEC20**a** *msh2*Δ*-1* | SAMN14647825 |
| 26. | YX61 | KN99α *msh2*Δ x JEC20**a** *msh2*Δ*-1* | SAMN14647826 |
| 27. | YX64 | KN99α *msh2*Δ x JEC20**a** *msh2*Δ*-1* | SAMN14647827 |
| 28. | YX65 | KN99α *msh2*Δ x JEC20**a** *msh2*Δ*-1* | SAMN14647828 |
| 29. | YX70 | KN99α *msh2*Δ x JEC20**a** *msh2*Δ*-1* | SAMN14647829 |
| 30. | SJP596 | H99α x KN99**a** progeny #1 | SAMN16690437 |
| 31. | SJP597 | H99α x KN99**a** progeny #2 | SAMN16690438 |
| 33. | SJP599 | H99α x KN99**a** progeny #4 | SAMN16690439 |
| 34. | SJP600 | H99α x KN99**a** progeny #5 | SAMN16690440 |
| 35. | SJP601 | KN99α *msh2*Δ x KN99**a** progeny #1 | SAMN16690441 |
| 36. | SJP602 | KN99α *msh2*Δ x KN99**a** progeny #2 | SAMN16690442 |
| 37. | SJP603 | KN99α *msh2*Δ x KN99**a** progeny #3 | SAMN16690443 |
| 38. | SJP604 | KN99α *msh2*Δ x KN99**a** progeny #4 | SAMN16690444 |
| 39. | SJP605 | KN99α *msh2*Δ x KN99**a** progeny #5 | SAMN16690445 |
| 40. | SJP606 | KN99α *msh2*Δ x KN99**a** *msh2*Δ*-1* progeny #1 | SAMN16690446 |
| 41. | SJP607 | KN99α *msh2*Δ x KN99**a** *msh2*Δ*-1* progeny #2 | SAMN16690447 |
| 42. | SJP608 | KN99α *msh2*Δ x KN99**a** *msh2*Δ*-1* progeny #3 | SAMN16690448 |
| 43. | SJP609 | KN99α *msh2*Δ x KN99**a** *msh2*Δ*-1* progeny #4 | SAMN16690449 |
| 44. | SJP610 | KN99α *msh2*Δ x KN99**a** *msh2*Δ*-1* progeny #5 | SAMN16690450 |
| 45. | SJP611 | KN99α *msh2*Δ x KN99**a** *msh2*Δ*-2* progeny #1 | SAMN16690451 |
| 46. | SJP612 | KN99α *msh2*Δ x KN99**a** *msh2*Δ*-2* progeny #2 | SAMN16690452 |
| 47. | SJP613 | KN99α *msh2*Δ x KN99**a** *msh2*Δ*-2* progeny #3 | SAMN16690453 |
| 48. | SJP614 | KN99α *msh2*Δ x KN99**a** *msh2*Δ*-2* progeny #4 | SAMN16690454 |
| 49. | SJP615 | KN99α *msh2*Δ x KN99**a** *msh2*Δ*-2* progeny #5 | SAMN16690455 |
| 50. | SJP616 | JEC21α x JEC20**a** progeny #1 | SAMN16690456 |
| 51. | SJP617 | JEC21α x JEC20**a** progeny #2 | SAMN16690457 |
| 52. | SJP618 | JEC21α x JEC20**a** progeny #3 | SAMN16690458 |
| 53. | SJP619 | JEC21α x JEC20**a** progeny #4 | SAMN16690459 |
| 55. | SJP621 | JEC21α x JEC20**a** *msh2*Δ*-1* progeny #1 | SAMN16690460 |
| 56. | SJP622 | JEC21α x JEC20**a** *msh2*Δ*-1* progeny #2 | SAMN16690461 |
| 57. | SJP623 | JEC21α x JEC20**a** *msh2*Δ*-1* progeny #3 | SAMN16690462 |
| 58. | SJP624 | JEC21α x JEC20**a** *msh2*Δ*-1* progeny #4 | SAMN16690463 |
| 59. | SJP625 | JEC21α x JEC20**a** *msh2*Δ*-1* progeny #5 | SAMN16690464 |
| 60. | SJP626 | JEC21α *msh2*Δ*-1* x JEC20**a** *msh2*Δ*-1* progeny #1 | SAMN16690465 |
| 61. | SJP627 | JEC21α *msh2*Δ*-1* x JEC20**a** *msh2*Δ*-1* progeny #2 | SAMN16690466 |
| 62. | SJP628 | JEC21α *msh2*Δ*-1* x JEC20**a** *msh2*Δ*-1* progeny #3 | SAMN16690467 |
| 63. | SJP629 | JEC21α *msh2*Δ*-1* x JEC20**a** *msh2*Δ*-1* progeny #4 | SAMN16690468 |
| 64. | SJP630 | JEC21α *msh2*Δ*-1* x JEC20**a** *msh2*Δ*-1* progeny #5 | SAMN16690469 |
| 65. | SJP631 | JEC21α *msh2*Δ*-2* x JEC20**a** *msh2*Δ*-1* progeny #1 | SAMN16690470 |
| 66. | SJP632 | JEC21α *msh2*Δ*-2* x JEC20**a** *msh2*Δ*-1* progeny #2 | SAMN16690471 |
| 67. | SJP633 | JEC21α *msh2*Δ*-2* x JEC20**a** *msh2*Δ*-1* progeny #3 | SAMN16690472 |
| 68. | SJP634 | JEC21α *msh2*Δ*-2* x JEC20**a** *msh2*Δ*-1* progeny #4 | SAMN16690473 |
| 69. | SJP635 | JEC21α *msh2*Δ*-2* x JEC20**a** *msh2*Δ*-1* progeny #5 | SAMN16690474 |
