## Supplementary figures and images for "Factors enforcing the species boundary between the human pathogens *Cryptococcus neoformans* and *Cryptococcus deneoformans*"

### S1 Figure

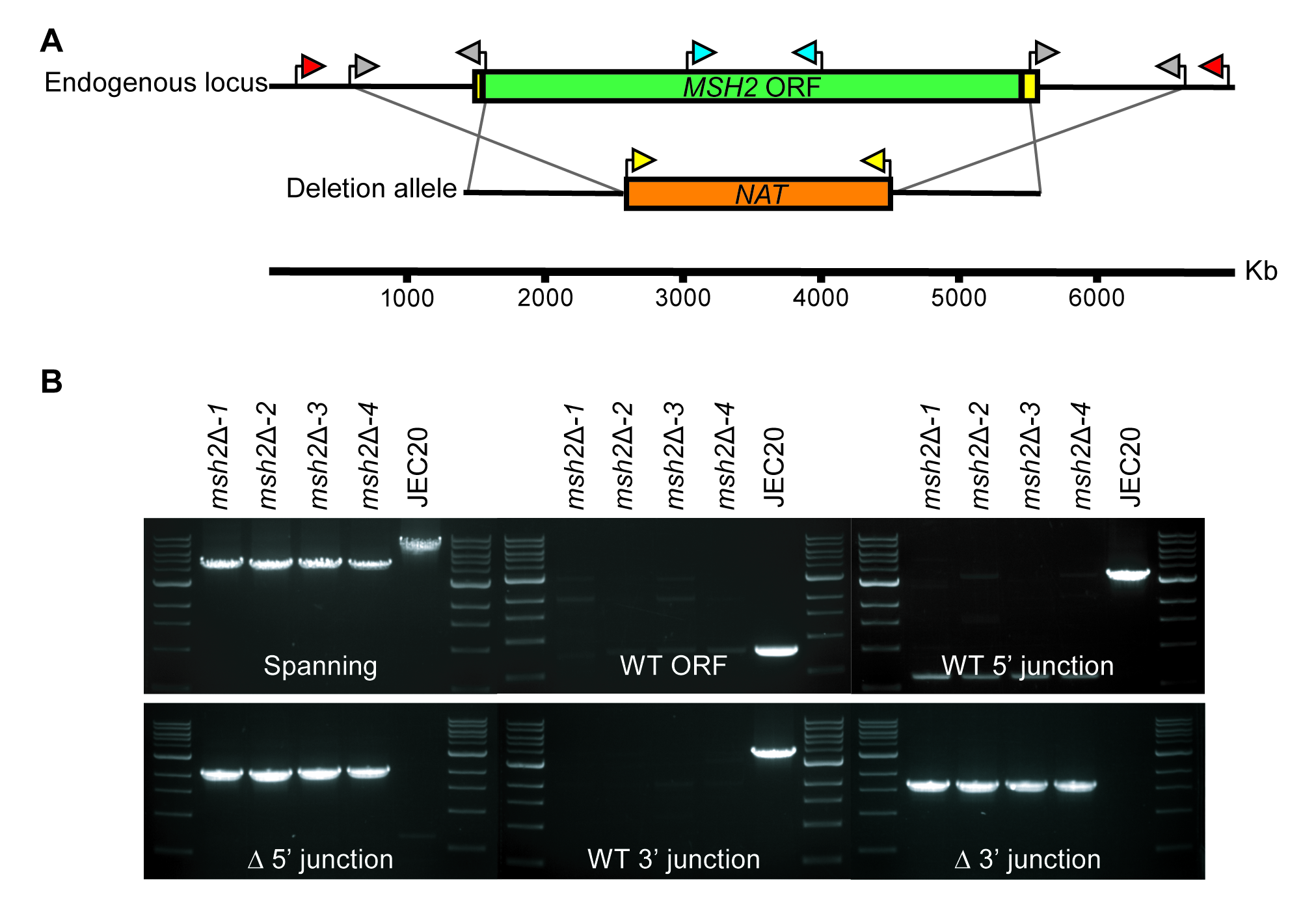

### S2 Figure

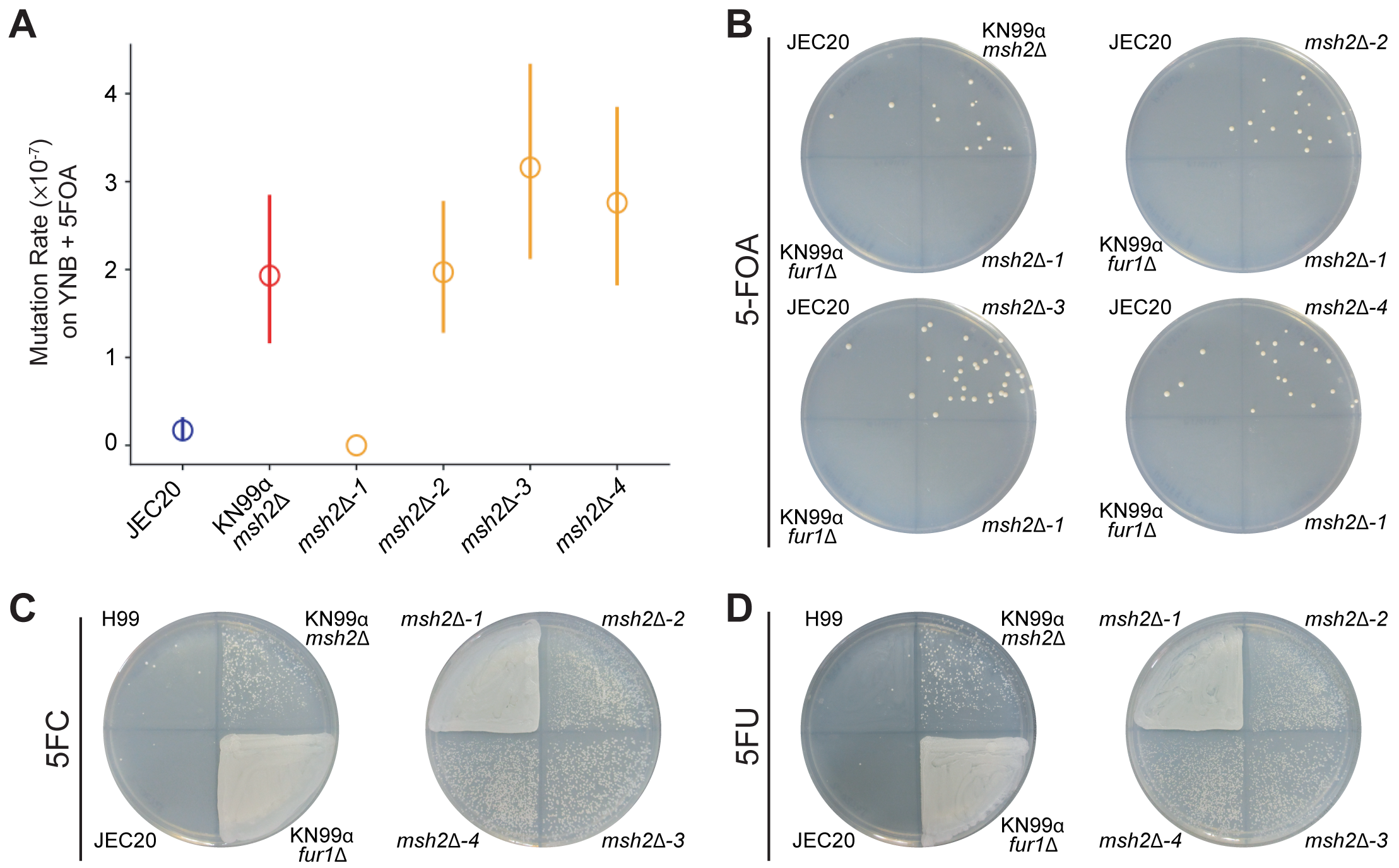

### S3 Figure

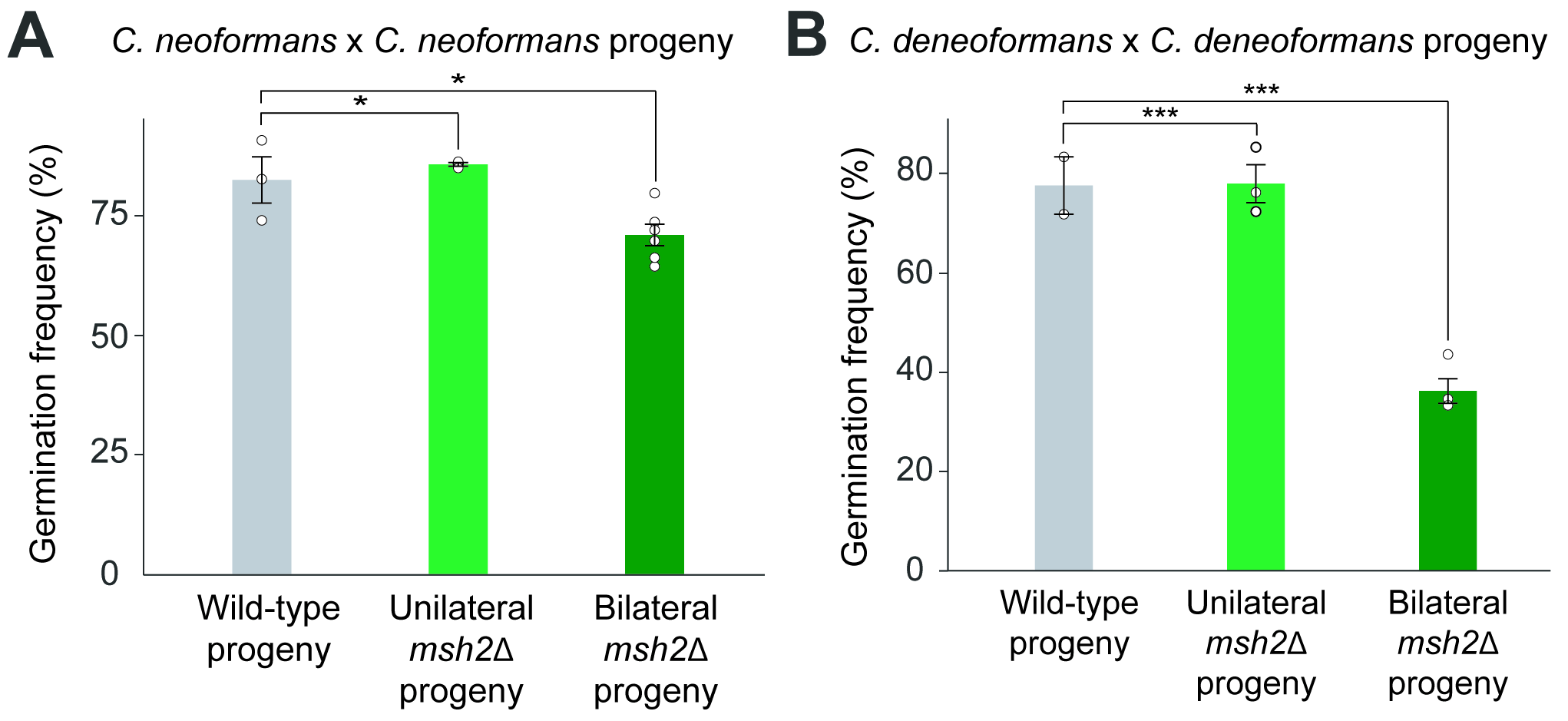

### S4 Figure

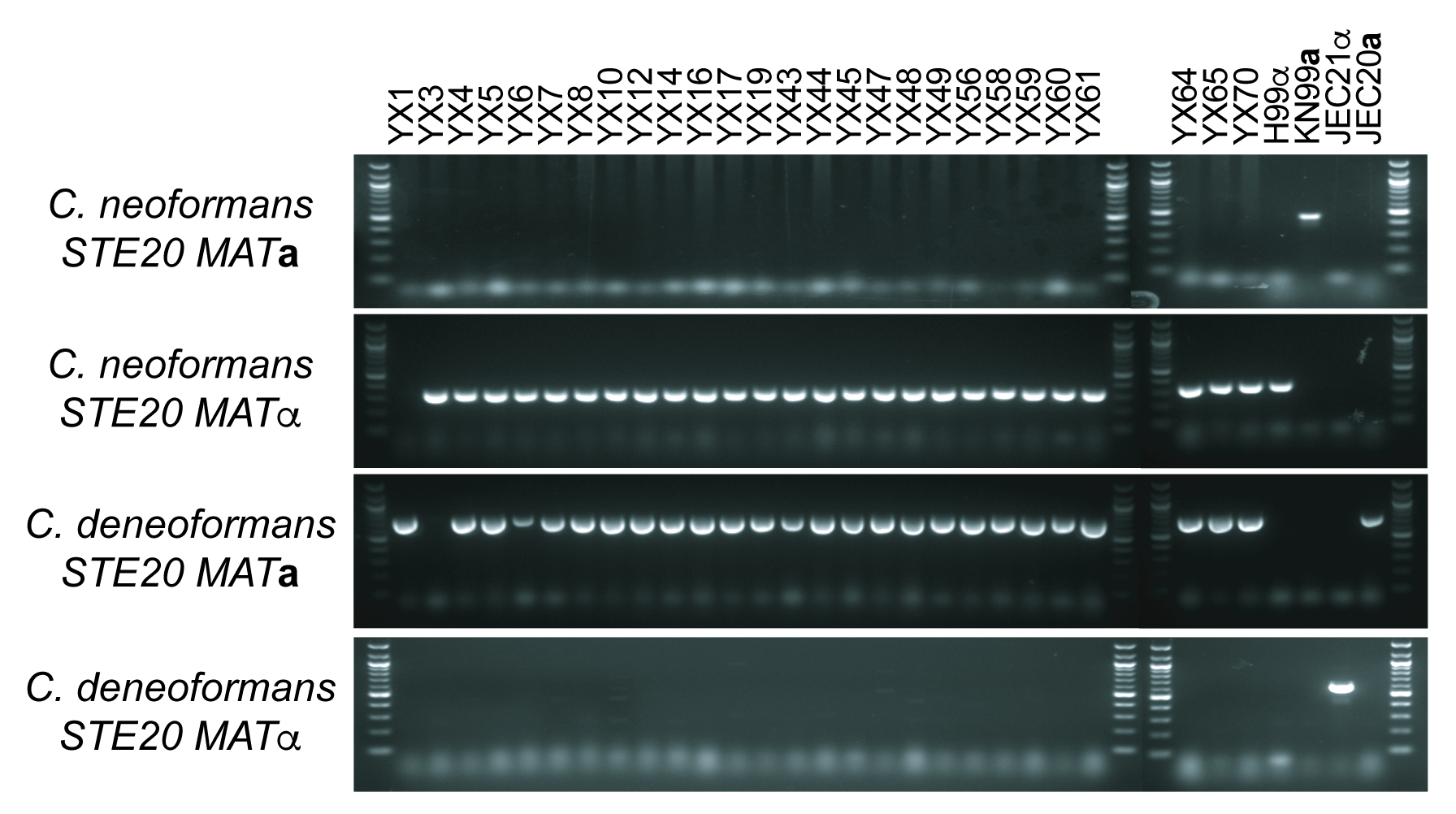

### S5 Figure

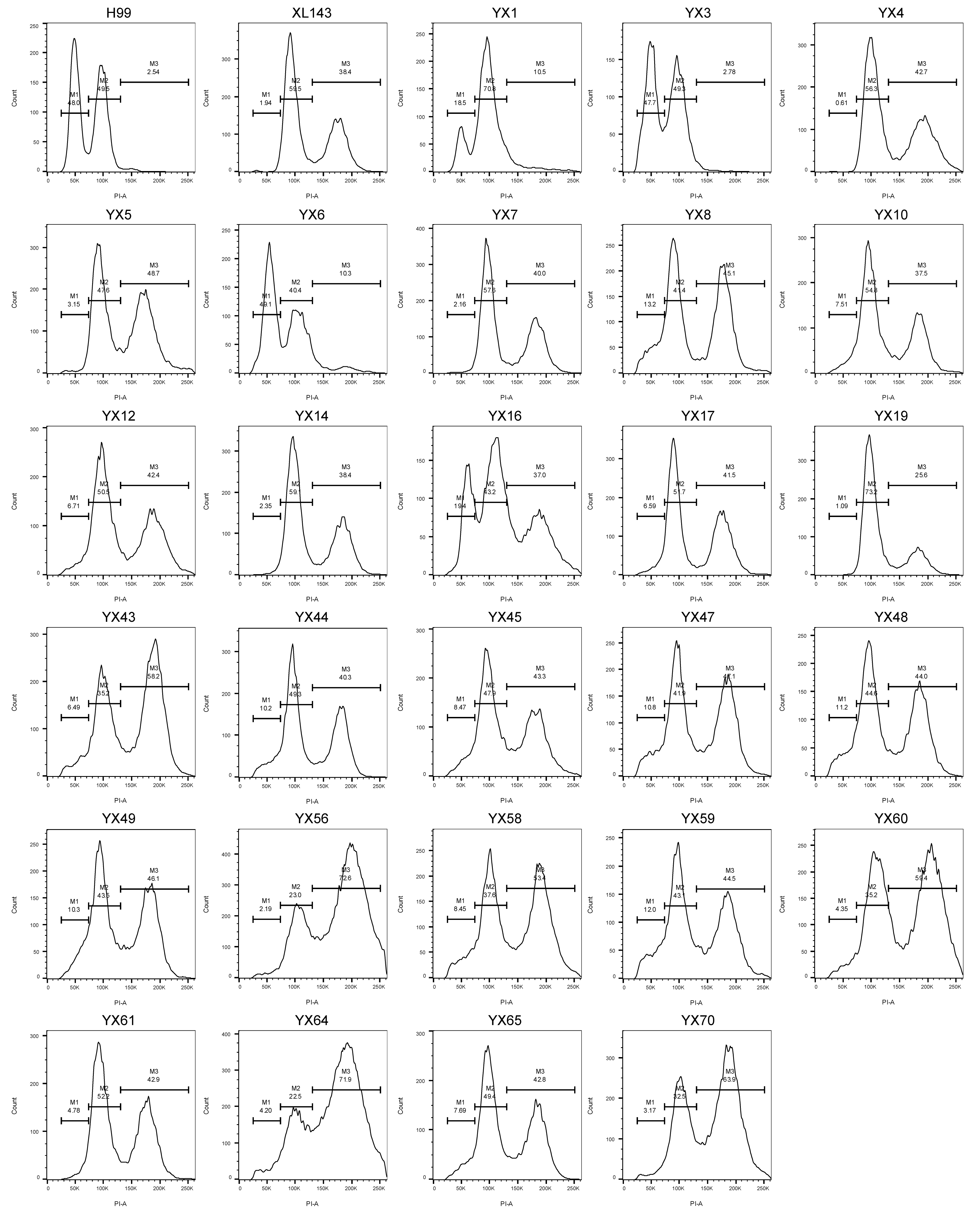

### S6 Figure

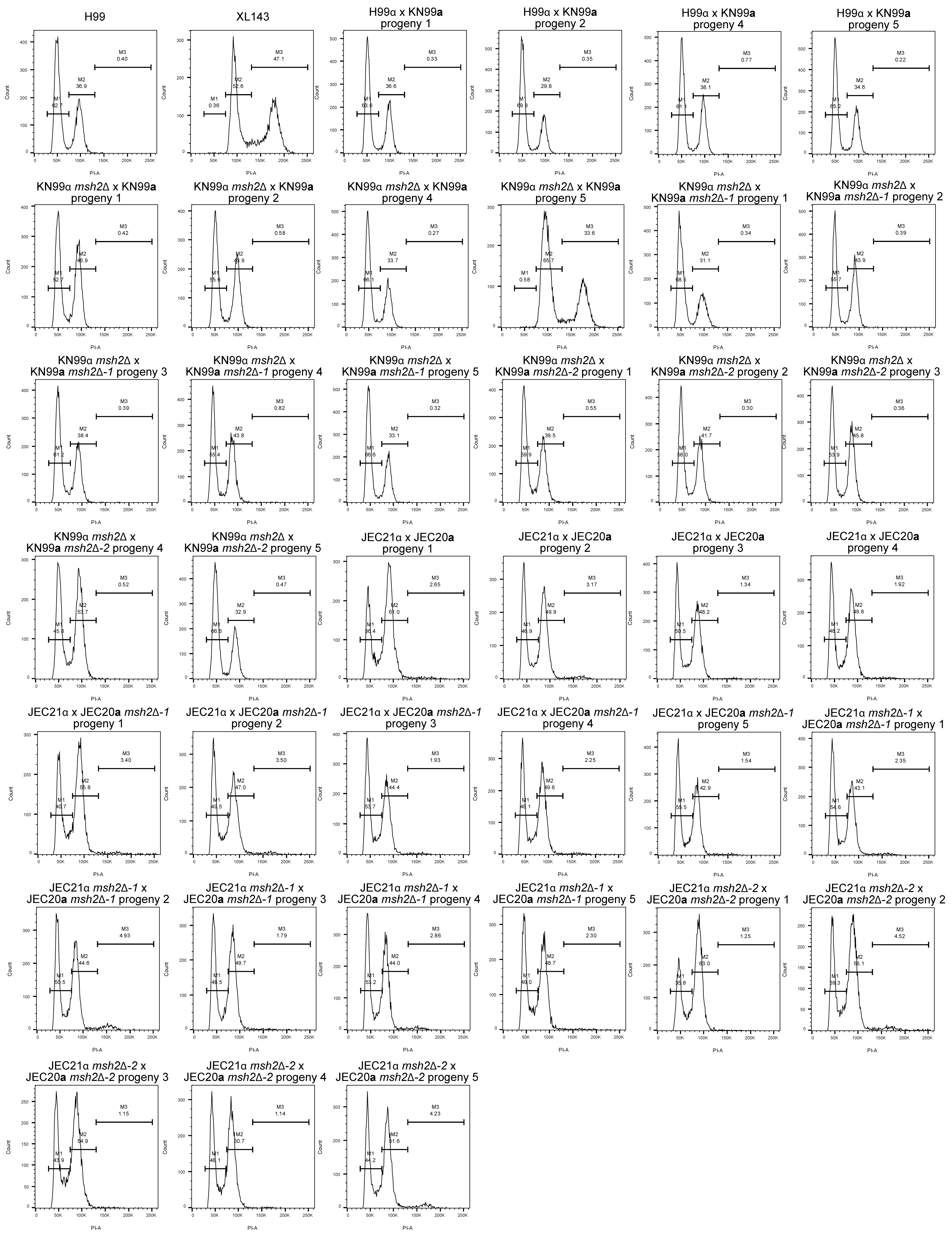

### S7 Figure

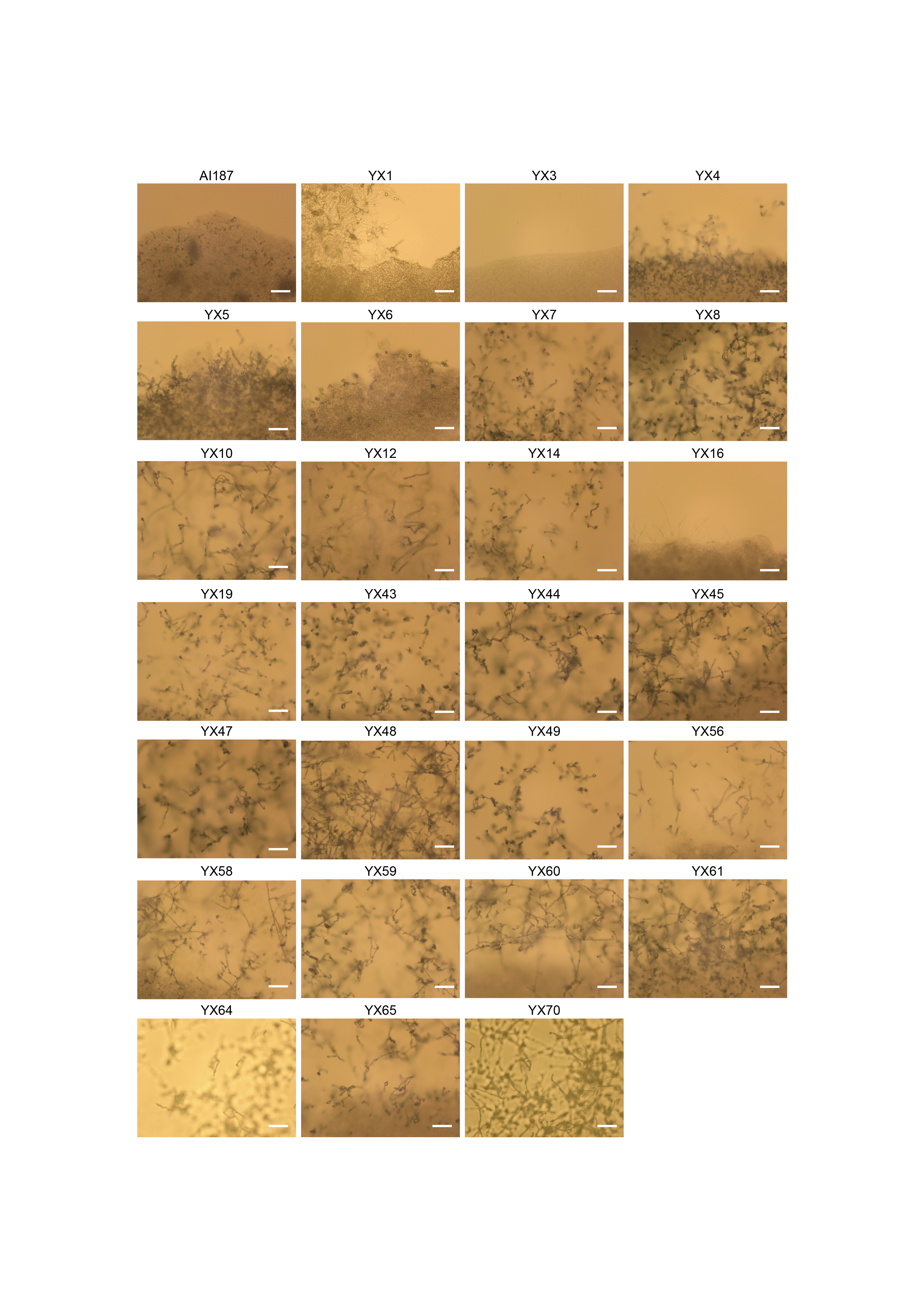

### S8 Figure

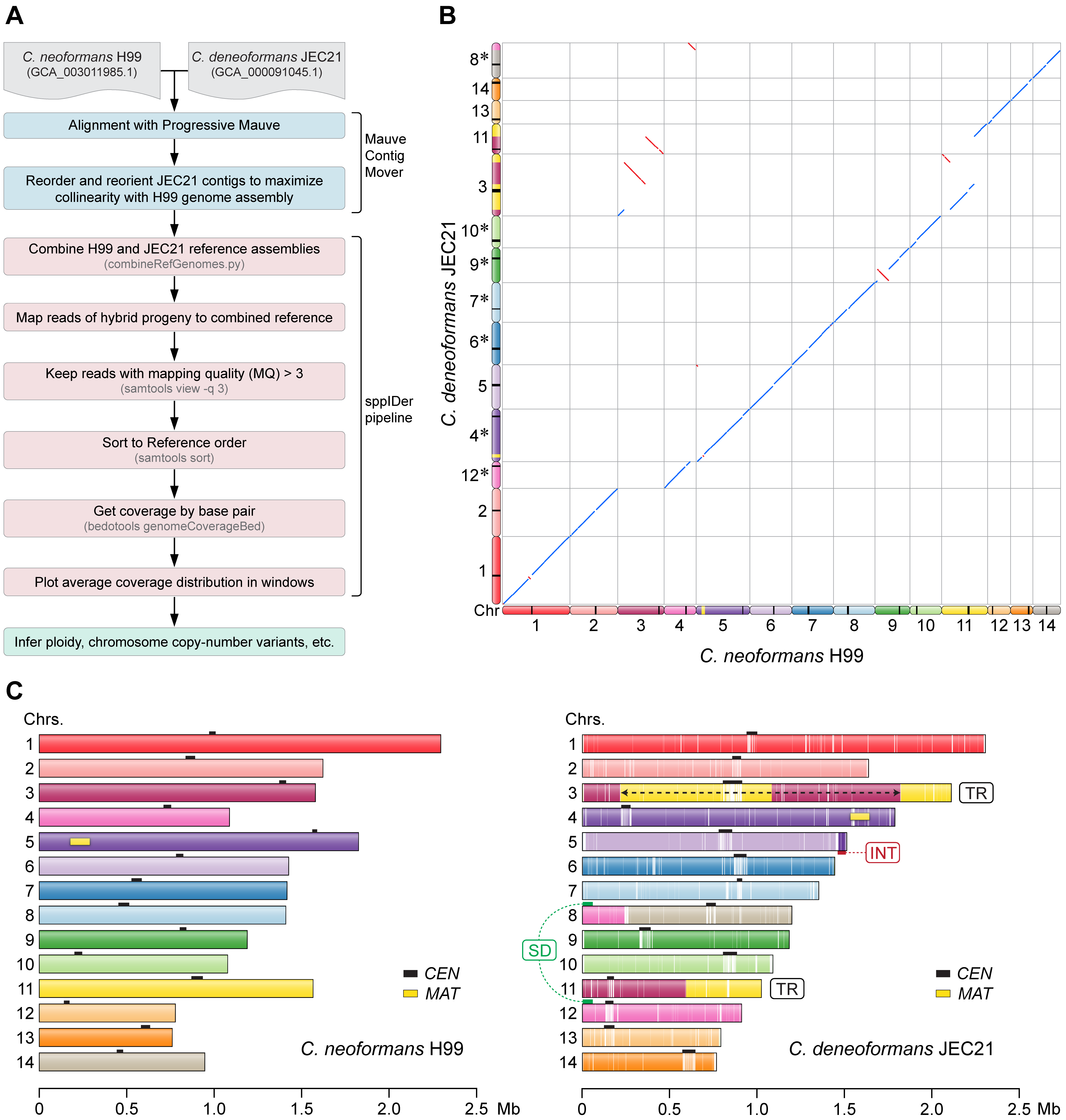

### S9 Figure

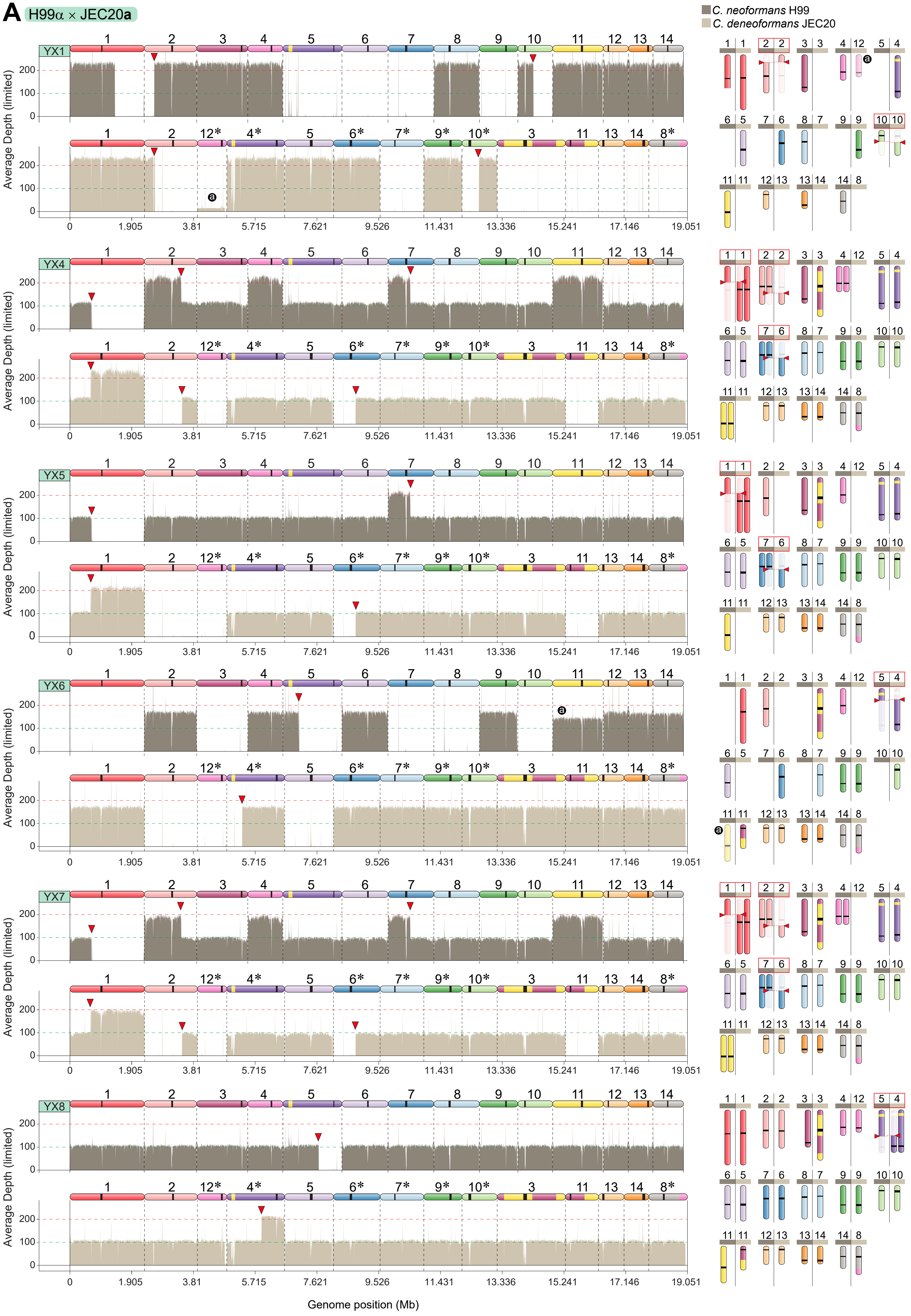

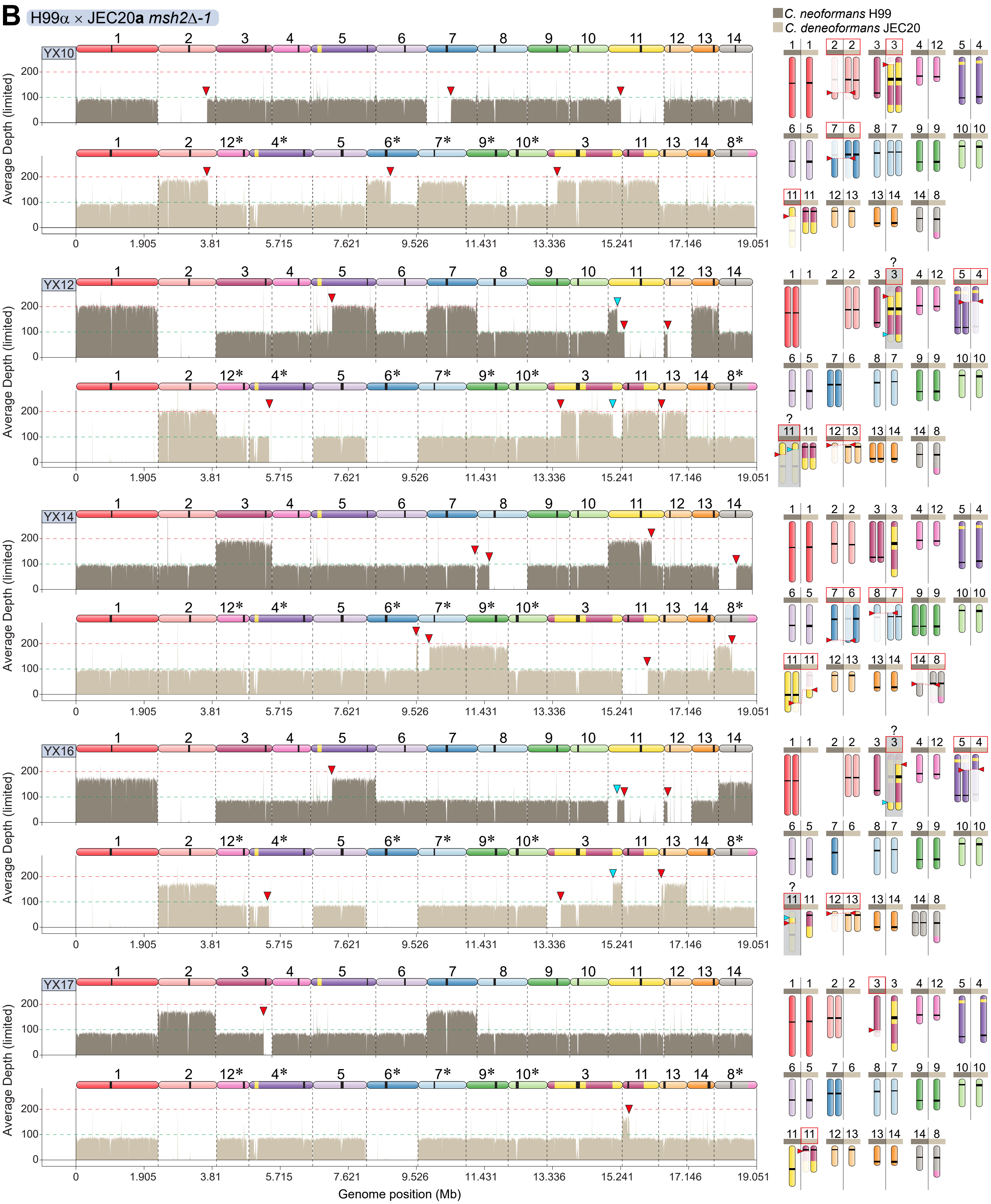

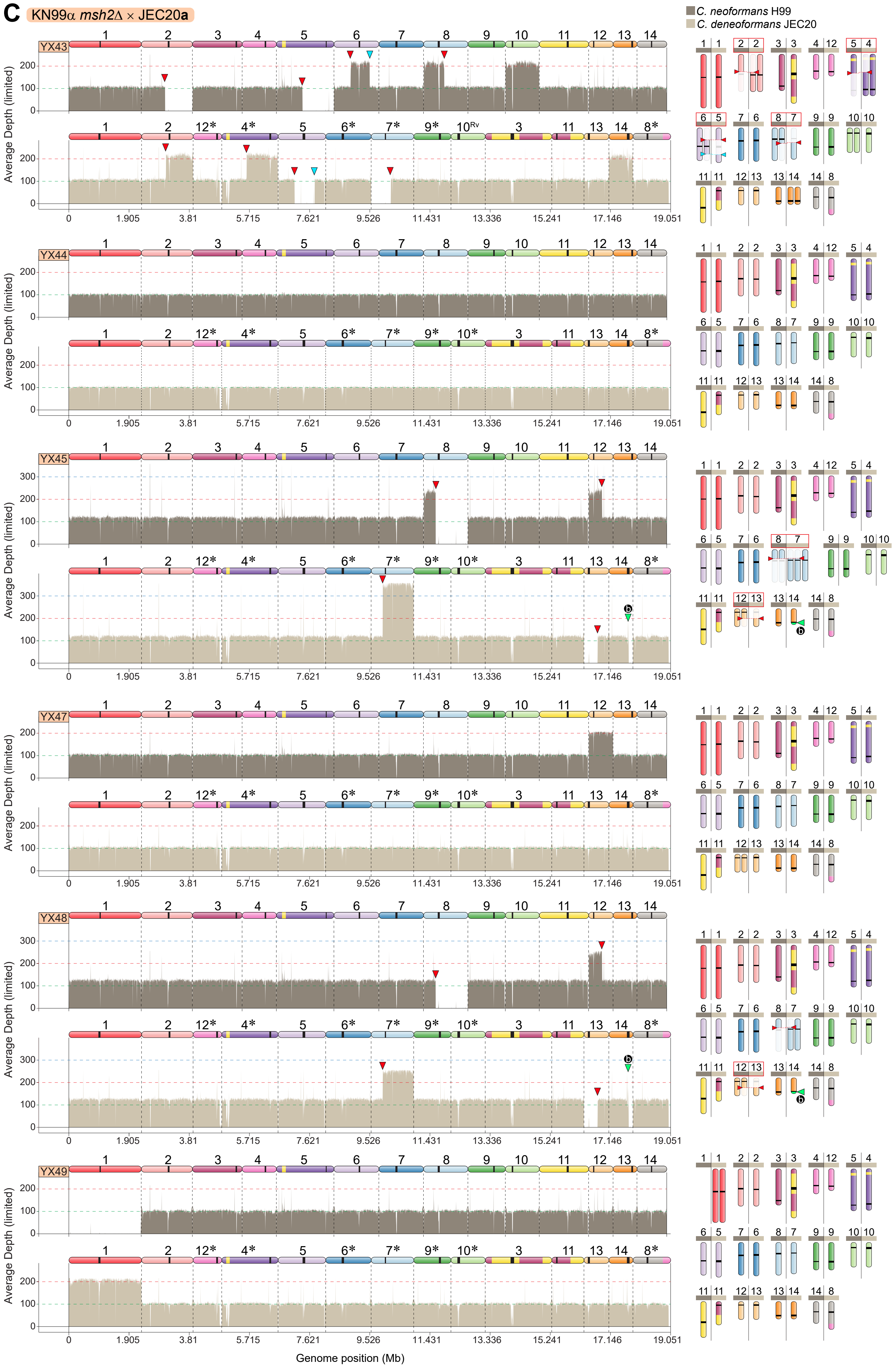

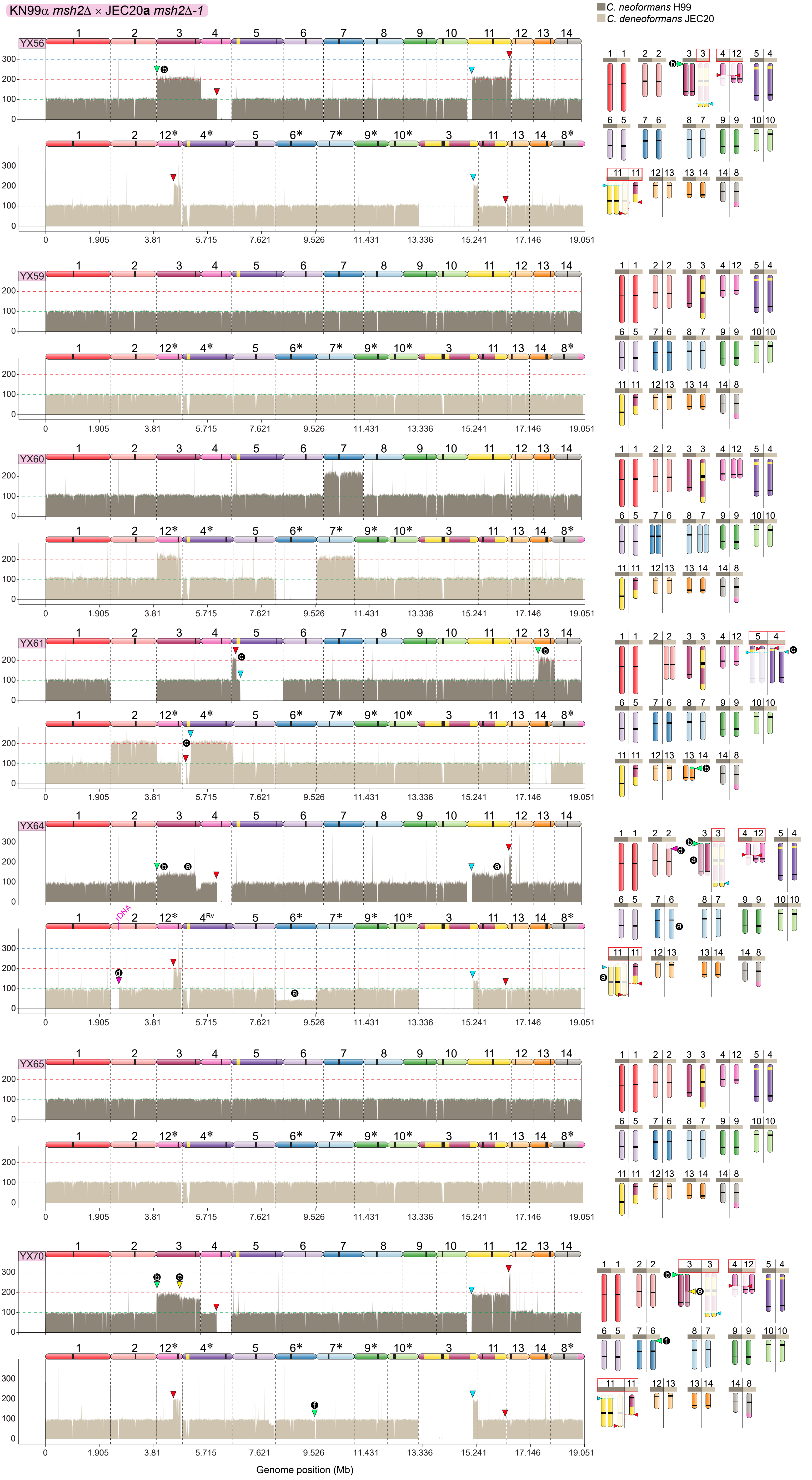

### S10 Figure

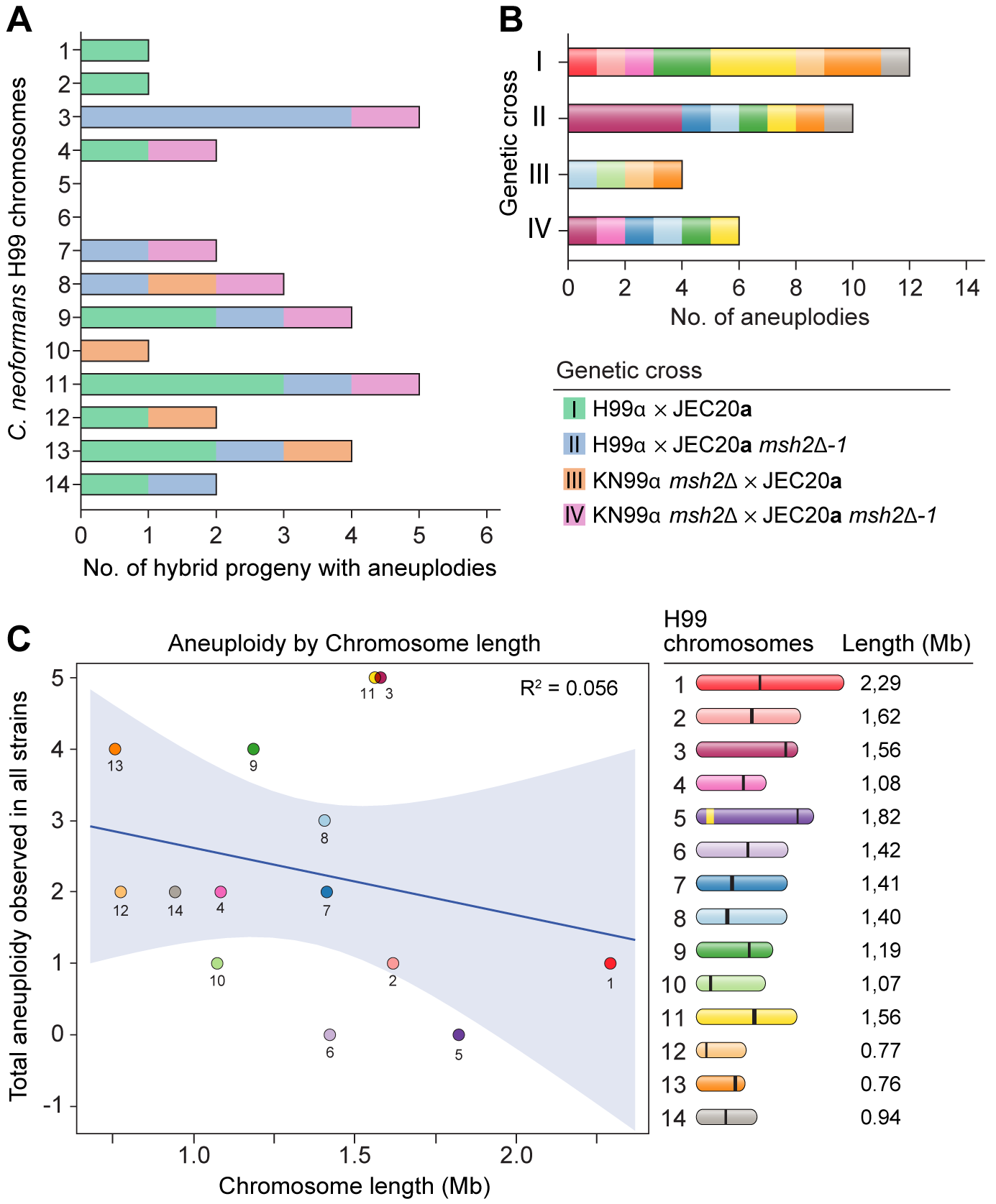

### S11 Figure

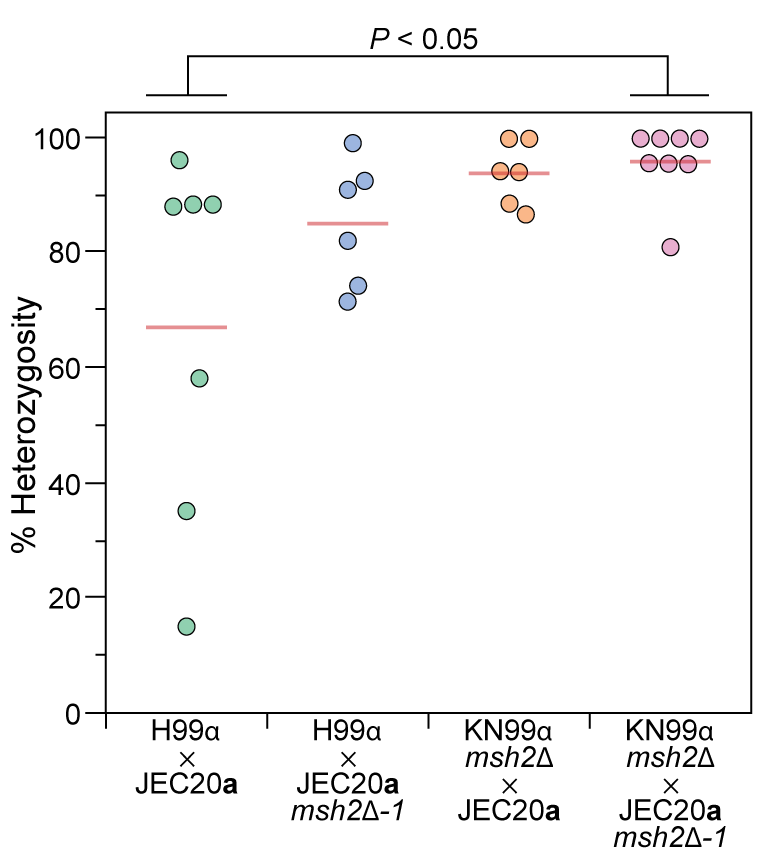

### S12 Figure

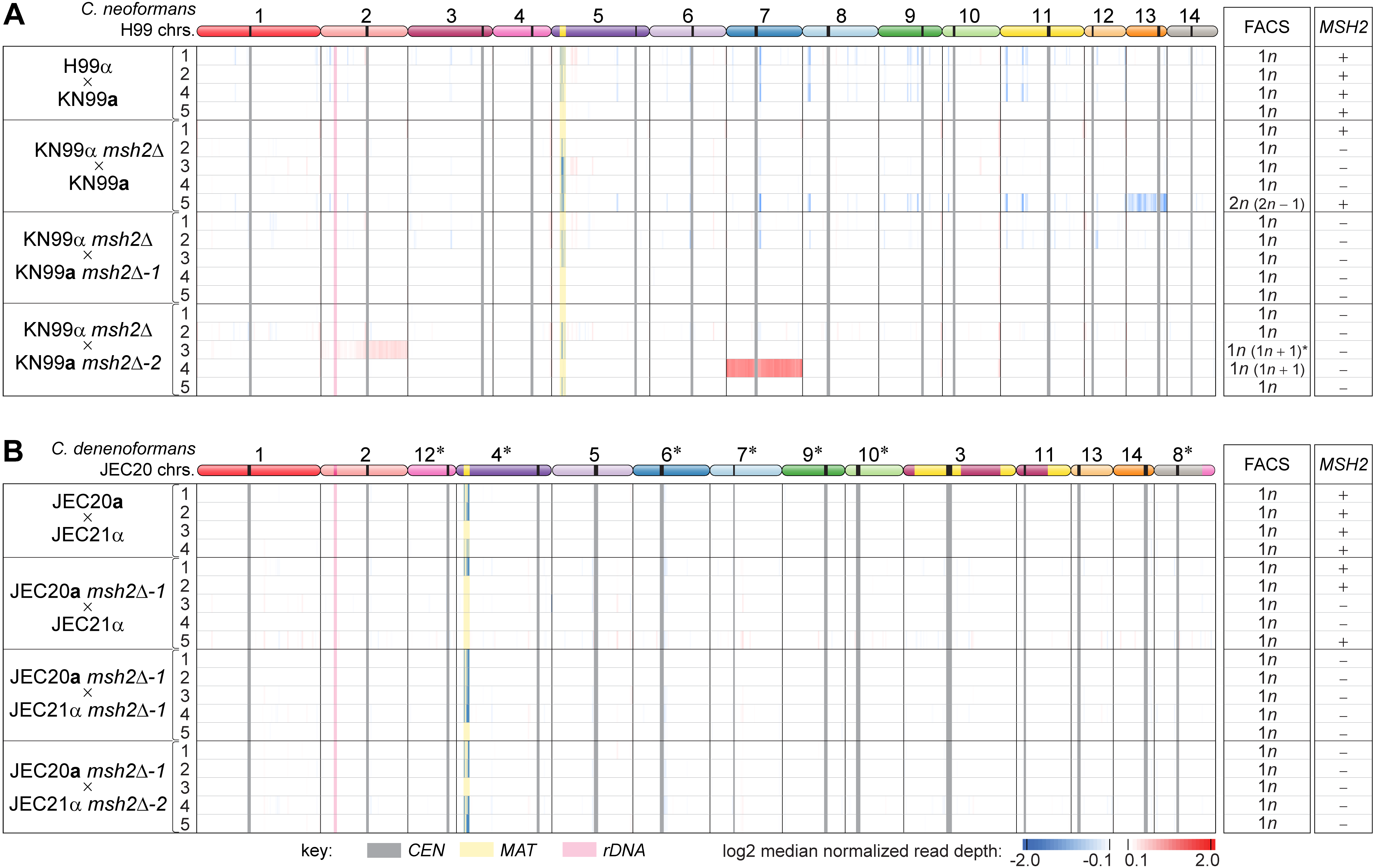

### S13 Figure

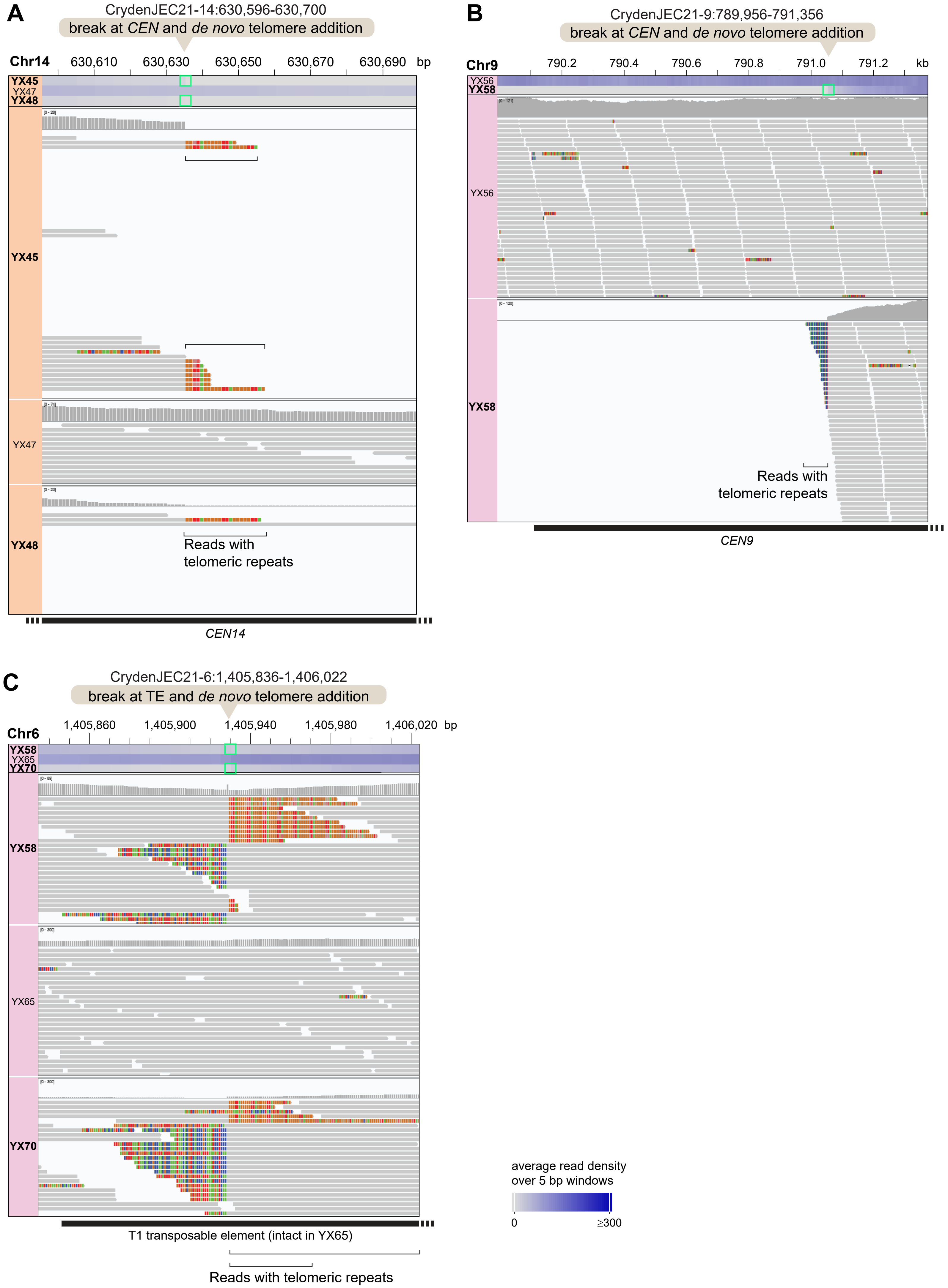

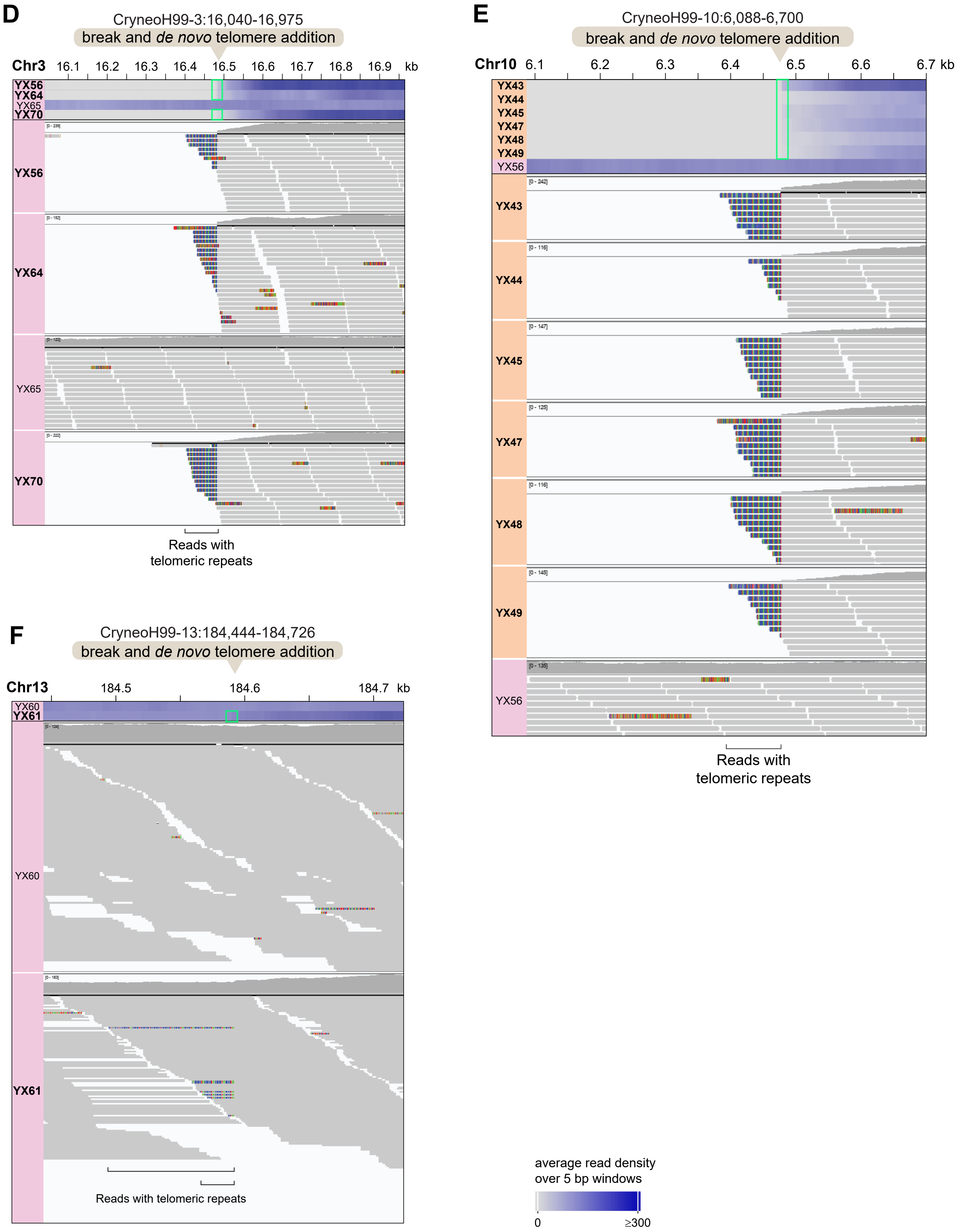

### S14 Figure

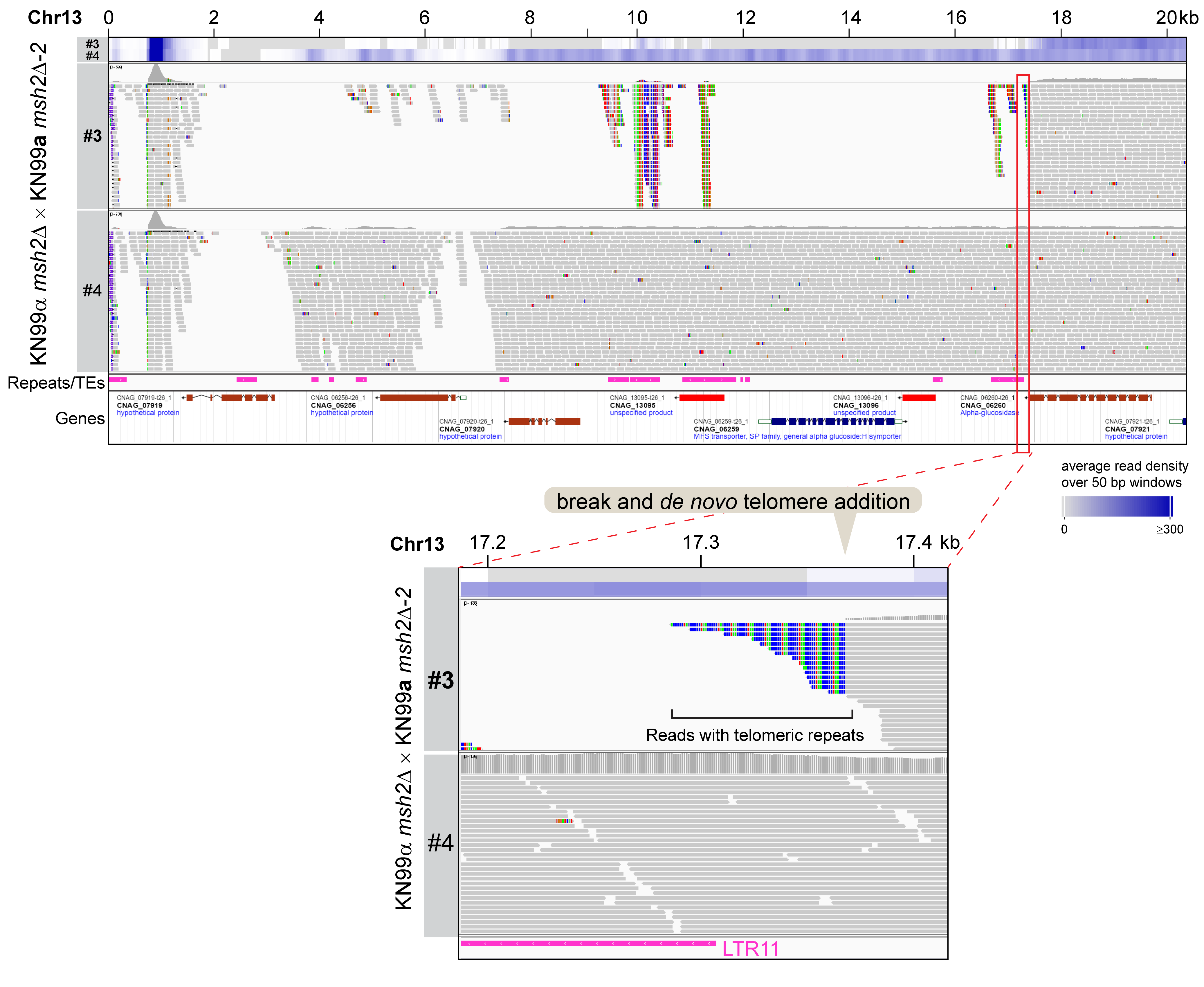
